# Persistent Epigenetic Reprogramming of Sweet Taste by Diet

**DOI:** 10.1101/2020.03.25.007773

**Authors:** Anoumid Vaziri, Morteza Khabiri, Brendan T. Genaw, Christina E. May, Peter L. Freddolino, Monica Dus

## Abstract

Diets rich in sugar, salt, and fat alter taste perception and food intake, leading to obesity and metabolic disorders, but the molecular mechanisms through which this occurs are unknown. Here we show that in response to a high sugar diet, the epigenetic regulator Polycomb Repressive Complex 2.1 (PRC2.1) persistently reprograms the sensory neurons of *D. melanogaster* flies to reduce sweet sensation and promote obesity. In animals fed high sugar, the binding of PRC2.1 to the chromatin of the sweet gustatory neurons is redistributed to repress a developmental transcriptional network that modulates the responsiveness of these cells to sweet stimuli, reducing sweet sensation. Importantly, half of these transcriptional changes persist despite returning the animals to a control diet, causing a permanent decrease in sweet taste. Our results uncover a new epigenetic mechanism that, in response to the dietary environment, regulates neural plasticity and feeding behavior to promote obesity.

## Introduction

Diets high in processed foods promote higher calorie intake and weight gain, increasing the risk for chronic and metabolic diseases (*1*). How these foods cause overconsumption, however, is still unclear. Processed foods are high in salt, and fat, which we are genetically programmed to like because of their high caloric-density (*2, 3*). Interestingly, evidence is emerging that the levels of salt, sugar, and fat in diets can alter taste sensation in humans (*4–7*), raising the question of whether these sensory changes may influence food intake, obesity, and metabolic disease (*8, 9*). This idea is supported by a number of recent animal studies which found changes in taste, neural responses, and food preferences in rodents fed high nutrient diets (*10–17*). However, due to the complexity of the mammalian taste system and the lack of genetic tools, we know next to nothing about the molecular mechanisms through which diet composition affects taste sensation and obesity. Thus, studies in genetically tractable model organisms could help shed light on this question and help us define evidence-based strategies to curb the prevalence of obesity and metabolic disease, which currently affects billions of people worldwide.

We recently found that high dietary sugar dulls the responses of the *D*. *melanogaster* taste neurons to sweet stimuli, causing higher food intake and weight gain, arguing that the effects of diet on taste are conserved (*18, 19*). In this manuscript we exploited the exquisite genetics tools of the fly and the relative simplicity of its sensory system to uncover the mechanisms through which high levels of dietary sugar reshape the sensory neurons to promote weight gain and obesity. We report that the Polycomb Repressive Complex 2.1 (PRC2.1), a chromatin silencing complex conserved from plants to humans (*20*), tunes the activity of the sweet sensory neurons and taste sensation in response to the food environment by repressing a transcriptional program that shapes the synaptic, signaling, and metabolic properties of these cells.

Interestingly, this diet-dependent transcriptional remodeling persisted even when animals were returned to the control diet, leading to lasting changes in sweet taste sensation that depended on the constitutive activity of PRC2.1. Together our findings suggest that diet composition activates epigenetic mechanisms that reprogram sensory responses to food; this sensory reprogramming determines the perception of future stimuli, leading to long-lasting alterations in behavior that increase the risk for obesity and metabolic disease.

## Results

### PRC2.1 modulates sweet taste in response to the food environment

*Drosophila melanogaster* flies fed high dietary sugar experience lower sweet taste sensation as a result of the decreased responsiveness of the sweet sensory neurons to sugar stimuli (*18*). Given the importance of sensory cues to control eating, and recent data that diet also impacts taste in mammals (*10–15, 18*), we set out to identify the molecular mechanisms through which the food environment shapes sensory responses. Since sweet taste deficits develop within 2-3 days upon exposure to the high sugar diet and independently of weight gain (*18*), we reasoned that gene regulatory mechanisms may be involved in modulating the responses of the sensory neurons. To test this hypothesis, we conducted a screen for gene regulatory and epigenetic factors necessary for the sweet taste defects caused by a high sugar diet. To do this, we fed control (*w1118_CS_*) and mutant flies a control diet (CD, ∼5% sucrose) or a diet supplemented with 30% sucrose (sugar diet, SD) for 7 days and then tested their ability to taste using the Proboscis Extension Response (*21*). This behavioral assay measures taste responses by quantifying the amount of proboscis extension (0= no extension, 0.5=partial extension, 1= full extension) when the fly labellum – where the dendrites and cell bodies of the taste neurons are located (Fig. S1A) – is stimulated with three different concentrations of sucrose (30%, 10%, 5%); this generates a taste curve where flies respond more intensely to higher sugar stimuli (Fig. 1B, gray circles). Flies fed a sugar diet show a marked decrease in PER compared to control diet flies (Fig. 1B, gray squares); however, mutants for the core Polycomb Repressive Complex 2 (PRC2) – which includes the histone 3 lysine 27 (H3K27) methyltransferase *Enhancer of Zeste* (*E(z)*), and the obligate accessory factors *Suppressor of zeste 12,* (*Su(z)12*) and *extra sex combs,* (*esc*) (Fig. 1A) – had largely the same proboscis extension response (PER) on a control and sugar diet (Fig. 1B, right, red shades). To confirm the role of PRC2 in taste sensation deficits, we supplemented the control and sugar diet with EED226, a PRC2 inhibitor (herein referred to as EEDi) that destabilizes the core complex by binding to the tri-methyl H3K27 (H3K27me3) binding pocket of EED (the homologue of *esc* in *M. musculus*) (*22*). While animals fed a sugar diet plus vehicle (10% DMSO) experienced lower PER, those fed a SD+EEDi retained normal sweet taste responses (Fig. 1C), consistent with results from the PRC2 mutants. Thus, mutations and inhibition of PRC2 prevents the blunting of sweet taste that occurs in the high sugar food environment.

**Fig. 1.**
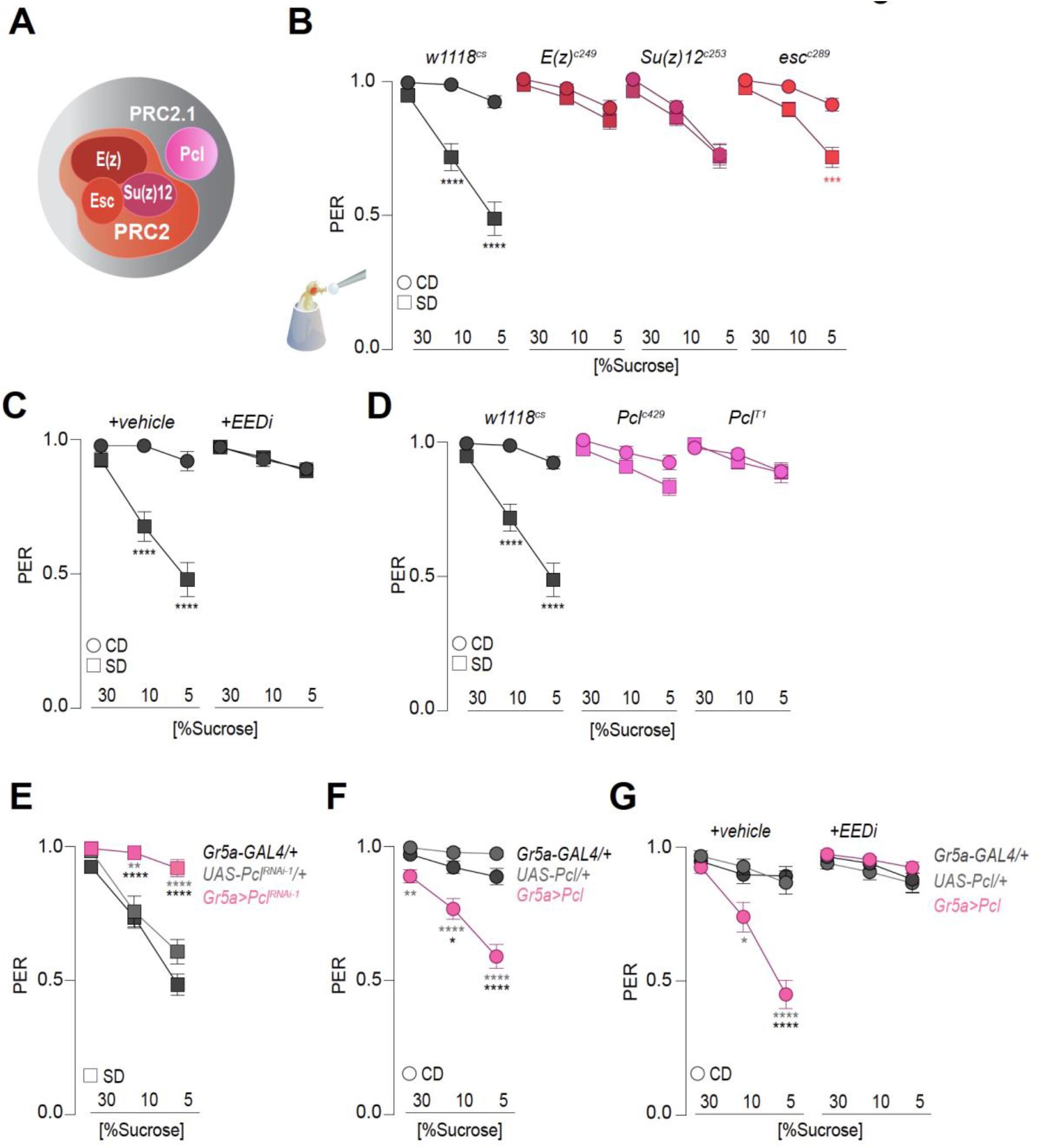
PRC2.1 modulates sweet taste in response to the food environment. **(A)** Schematic of *Drosophila melanogaster* PRC2 core complex consisting of *Enhancer of zeste* (*E(z))*, *Suppressor of zeste* (*Su(z)12)*, *extra sex combs (esc*) and the accessory protein *Polycomb-like* (*Pcl*). **(B-G)** Taste responses (y axis) to stimulation of the labellum with 30%, 10% and 5% sucrose (x axis) of age-matched males (**B)** *w1118_cs_* (gray), *E(z)_c249/+_* (dark red), *Su(z)12_c253/+_* (fuschia), and *esc_c289/+_* (red) flies on a control (circle) or sugar (square) diet for 7 days. n = 34-68; **(C)** *w1118_cs_* flies on a control (circle) or sugar (square) diet supplemented with vehicle (10% DMSO) or 8 *μ*M EEDi for 7 days. n = 32-43; **(D)** *w1118_cs_* (gray), *Pcl_c429/+_* (pink) and *Pcl_T1/+_* (pink) flies on a control (circle) or sugar (square) diet for 7 days. n = 36-82; **(E)** *Gr5a>Pcl_RNAi-1_* (pink), and parental transgenic controls (gray, crossed to *w1118_cs_*) flies on a sugar diet for 7 days. n = 42-63; **(F)** *Gr5a>Pcl* (pink), and parental transgenic controls (gray, crossed to *w1118_cs_*) flies on a control diet for 7 days. n = 36-61; **(G)** *Gr5a>Pcl* flies on a control diet supplemented with vehicle (10% DMSO) or 8 *μ*M EEDi for 7 days. n = 30-35. Data are shown as mean ± SEM, Kruskal-Wallis Dunn’s multiple comparisons, comparisons to control diet within genotype groups. **** *p* < 0.0001, *** *p* < 0.001, ** *p* < 0.01, \**p*< 0.05 for all panels in this figure.

In flies PRC2 forms two main subcomplexes, PRC2.1 and PRC2.2, which contain distinct accessory factors that can influence the targeting of the core complex to the chromatin (*23*). Mutations in the *Polycomb-like* (*Pcl*) gene, the accessory factor to PRC2.1, phenocopied PRC2 mutants and prevented sweet taste deficits in flies fed a sugar diet (Fig. 1D). In contrast, flies with deficits in the PRC2.2-members *Jumonji, AT rich interactive domain 2* (*Jarid2*) and *jing* still showed a blunting of sweet taste responses in flies fed a sugar diet (Fig. S1B). Interestingly, members of the Polycomb Repressive Complex 1 (PRC1) and the recruiter complex PhoRC were also not required for the taste changes in responses to a sugar diet (Fig. S1C and Fig. S1D-E). Thus, the PRC2.1 complex is necessary for the sensory changes that occur in the high sugar environment.

We next asked if PRC2.1 is required specifically in the sweet sensory neurons to decrease their responses to sweet stimuli on the sugar diet. To do this, we used the GAL4/UAS system to knock down *Pcl* in the sweet taste neurons using the *Gustatory receptor 5a* GAL4 driver*, Gr5a-GAL4,* which labels ∼60 cells in the proboscis of adult flies (24); we selected *Pcl* to narrow the effect to the PRC2.1 complex. Knockdown of *Pcl* in *Gr5a+* neurons using two independent RNAi transgenes (50% knockdown efficiency, Fig. S2A) prevented sweet taste deficits in animals fed a sugar diet (Fig. 1E, and Fig. S2B). *Pcl* knockdown, however, had no effect on sweet taste on a control diet (Fig. S2C), in accordance with the observation that *E(z)* and *Pcl* mutants have no effect on taste on a control diet (Fig. 1B, C) and suggesting that these phenotypes are uncovered only by the high sugar food environment.

Since *Pcl* is thought to target the PRC2 core complex to chromatin (*23*), we hypothesized that its overexpression may be sufficient to induce sweet taste deficits even in the absence of a high sugar food environment. Indeed, overexpression of *Pcl* specifically in the *Gr5a+* neurons induced sweet taste deficits in flies fed a control diet compared to transgenic controls (Fig. 1F). The effects of *Pcl* overexpression were abolished by treatment with the PRC2 inhibitor EEDi (Fig. 1G), arguing that *Pcl* causes sweet taste deficits entirely through the action of PRC2 and not through some yet unidentified mechanism. Importantly, *Pcl* overexpression had no effect on the number of *Gr5a+* neurons in the proboscis (Fig. S2D), and so the taste deficits cannot be attributed to a decrease in the number of cells. To exclude the possibility that the effects of manipulating *Pcl* on sweet taste were developmental, we used the temperature sensitive *tubulin-GAL80_ts_* transgene to limit expression of *UAS-Pcl* and *Pcl_RNAi-1_* only in adult flies. Switching the flies to the non-permissive temperature and diet 4 days post eclosion, resulted in the same effects on sweet taste as using the *Gr5a-GAL4* alone (Fig. S2E). Together, these experiments establish that the PRC2.1 complex is required cell-autonomously in the *Gr5a+* neurons to mediate the effects of a high sugar diet on sweet taste.

### *Pcl* mutant flies have normal sensory responses and are resistant to diet-induced obesity

Flies on a high sugar diet have lower sweet taste because the neural responses of the taste neurons to sweet stimuli are decreased (*18*). Since *Pcl* mutants have identical taste on a control and sugar diet, we hypothesized that the responses of the sensory neurons to sucrose stimulation should also be similar. To test this, we expressed the genetically encoded presynaptic calcium indicator *UAS*-*GCaMP6s-Brp-mCherry* (*25*) in the sweet sensing neurons and measured their *in vivo* responses to stimulation of the proboscis with 20% sucrose in *Pcl* mutant animals (Fig. 2A). Indeed, the responses to sucrose stimulation were identical in *Pcl* mutant flies fed a control diet and sugar diet (Fig. 2B), matching the behavioral data (Fig. 1); importantly, this rescue was not due to an increase in the number of sweet taste cells (Fig. 2C).

**Fig. 2.**
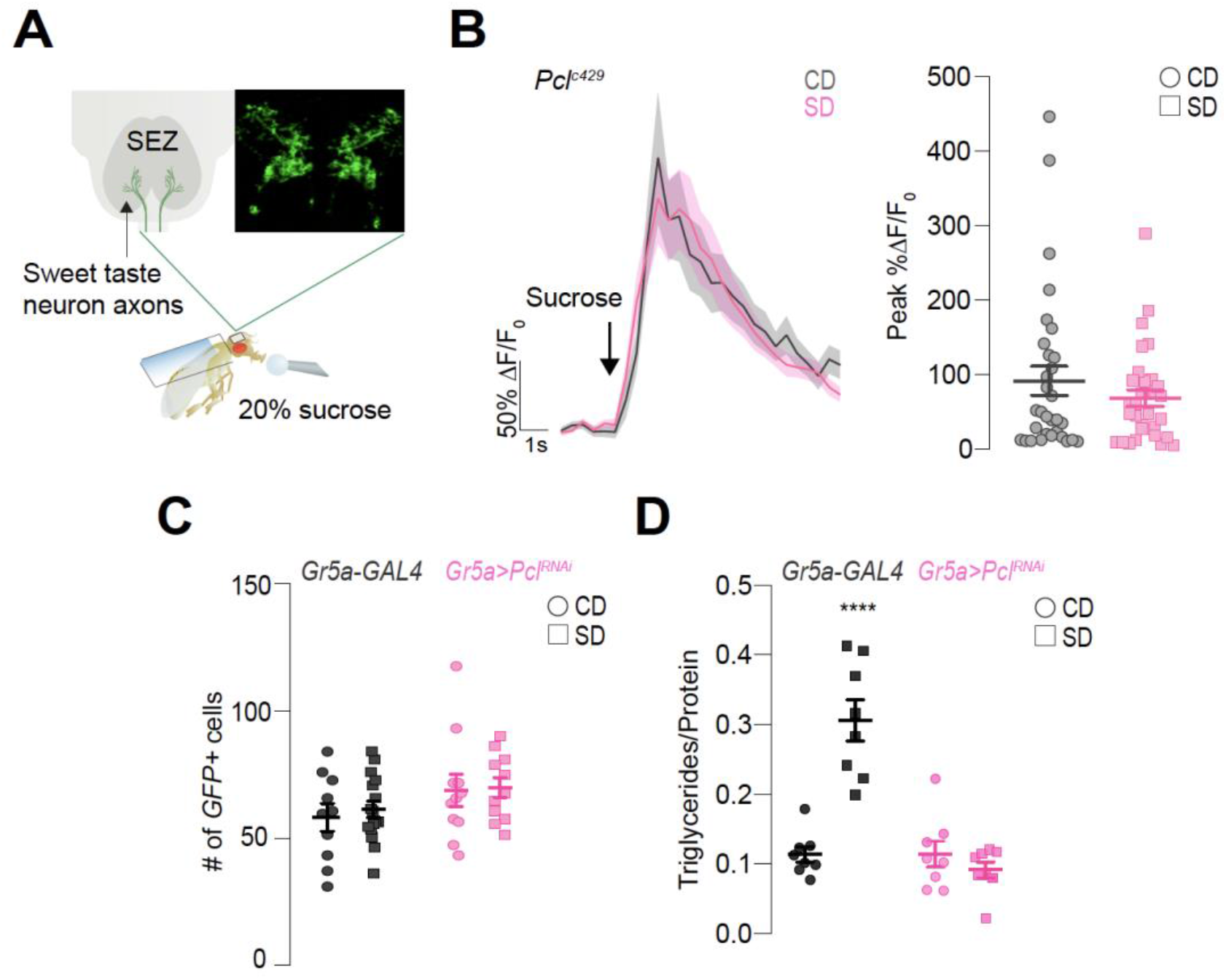
*Pcl* mutant flies have normal sensory responses and are resistant to diet-induced obesity. **(A)** Schematic showing the setup for *in vivo* calcium imaging: the fly proboscis is stimulated with 20% sucrose while recording from the <u>s</u>ub<u>e</u>sophageal <u>z</u>one (SEZ) which contains the presynaptic terminals of the sweet taste neurons (shown here labeled with *synaptotagmin::GFP*). **(B)** Average ΔF/F_0_ calcium response traces (left panel) and peak % ΔF/F_0_ (right panel) responses to sucrose stimulation of the proboscis (arrow) in age-matched male *Gr64f>GCaMP6s-Bruchpilot-mCherry;Pcl_c429_* flies fed a control (gray, circle) or sugar (pink, square) diet for 7 days, n = 32-34. **(C)** Quantification of GFP-labeled cells in the labella of *Gr64f;CD8-GFP* flies crossed to *w1118_cs_* (as control, dark gray) or *Gr64f;CD8-GFP>Pcl_RNAi_* (pink) on a control (circle) or sugar (square) diet for 7 days. n=5-16 flies, Kruskal-Wallis Dunn’s multiple comparisons, comparison to control diet of each genotype, no significance. **(D)** Triglyceride levels normalized to protein in *Gr5a>Pcl_RNAi_* (pink) and parental transgenic control flies (dark gray, crossed to *w1118_cs_*) fed a control (circle) or sugar (square) diet for 7 days. n=8, two-way ANOVA with Sidak’s multiple comparisons test, comparisons to control diet of each genotype. All data are shown as mean ± SEM, ∗∗∗∗*p* < 0.0001, ∗∗∗*p* < 0.001, ∗∗*p* < 0.01, and ∗*p* < 0.05 for all panels unless indicated.

We previously showed that restoration of sweet taste neuron activity in flies fed high dietary sugar protected them from diet-induced obesity (*18*). Since *Pcl* mutants abolished the deficits in neural and behavioral responses to sweetness in animals fed a high sugar diet, we anticipated that they should also prevent a diet-dependent increase in triglycerides. Indeed, sugar-diet flies with knockdown of *Pcl* in the *Gr5a+* neurons remained as lean as animals on a control diet (Fig. 2D), while triglycerides increased in control flies fed a sugar diet (Fig. 2D). Importantly, there was no difference in the levels of triglycerides between control and *Pcl* knockdown flies fed a control diet (Fig. 2D). Together, these data suggest that, in response to the food environment, *Pcl* modulates the responsiveness of the sweet gustatory neurons to promote diet-induced obesity.

### *Pcl* chromatin occupancy is redistributed in the high sugar environment

Our experiments show that PRC2.1 plays a critical role in the neural activity, behavior, and the metabolic state of animals exposed to the high sugar food environment. To identify the molecular mechanisms underlying these phenotypes, we measured the chromatin occupancy of *Pcl* in the ∼60 *Gr5a+* neurons using Targeted DNA Adenine Methyltransferase Identification (TaDa) (Fig. 3A) (*26–28*). To do this we generated *Dam::Pcl* (*UAS-LT3-Dam::Pcl*) transgenic flies and compared them to *Dam*-only flies (*UAS-LT3-Dam*) to control for non-specific methylation by the freely diffusing *Dam* protein (*26*) (Fig. 3A) and to obtain a measure of chromatin accessibility *in vivo* (CATaDa) (*29*). To specifically profile *Pcl* binding to chromatin in the sweet sensory neurons and limit the induction of *Dam*, we expressed the *Dam::Pcl* and *Dam* transgenes in combination with *Gr5a-GAL4*;*tubulin-GAL80_ts_*. To induce the expression of each UAS transgene we shifted the flies to the permissive temperature (28°C) for 20 hours after they had been exposed to a control or sugar diet for 3 days (Fig. 3A). We selected this time point because we previously showed that sweet taste defects developed within 3 days of exposure to the sugar diet (*18*).

**Fig. 3.**
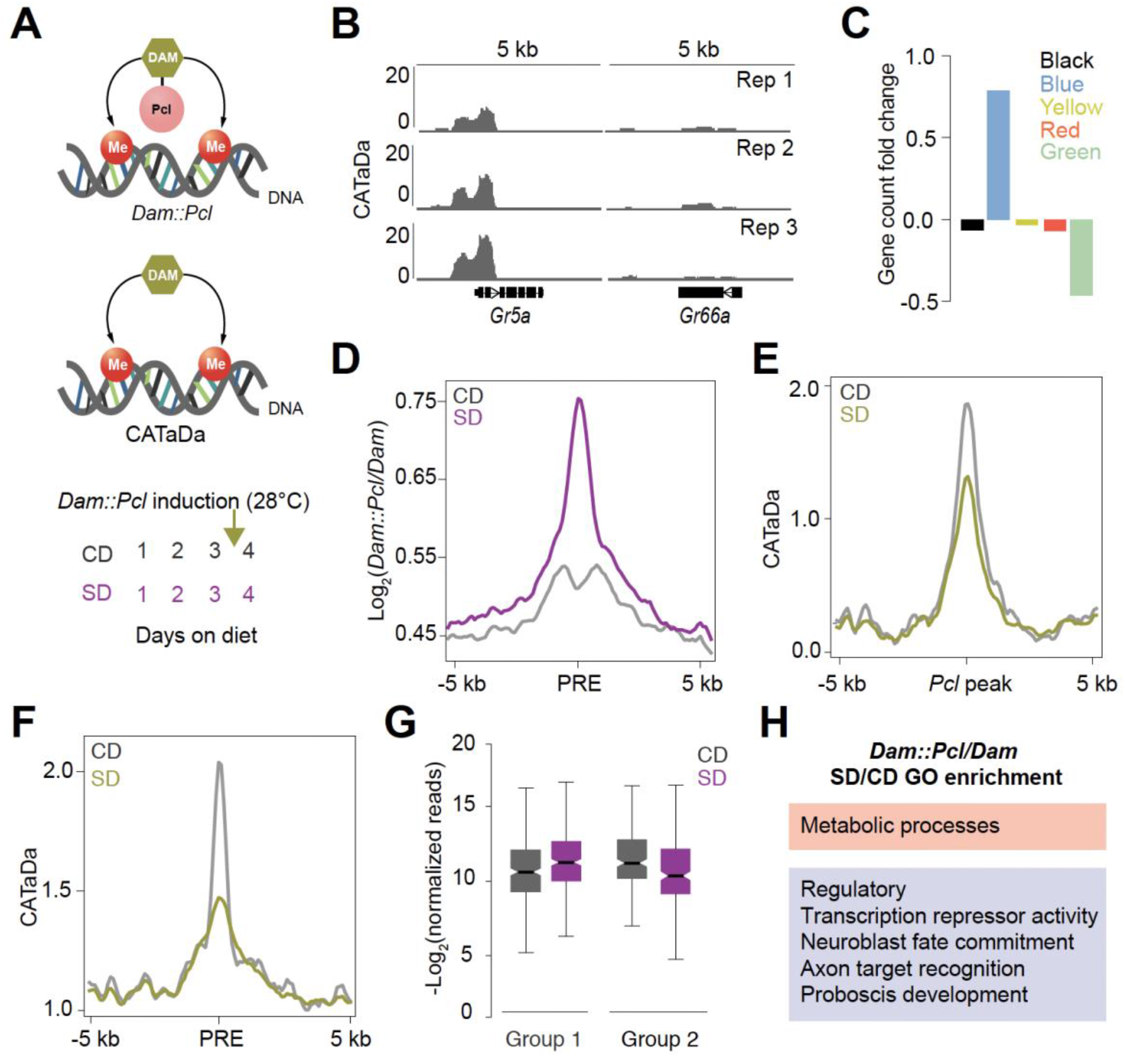
*Pcl* chromatin occupancy is redistributed in the high sugar environment. **(A)** Schematic of Targeted Dam-ID (TaDa) of *Dam::Pcl* and *Dam* (CATaDa, top panel) and of TaDa induction paradigm (bottom panel). Age-matched *Gr5a;tubulin-GAL80>UAS-LT3-Dam::Pcl* and *Gr5a;tubulin-GAL80>UAS-LT3-Dam* flies were placed on a control (gray) or sugar (purple) diet for 3 days at 20-21°C, then switched to 28°C between day 3-4 to induce *Dam* expression (arrow) (bottom panel). **(B)** CATaDa chromatin coverage (chromatin accessibility) normalized to 1x dm6 genome (reads per genome coverage) of the three biological replicates from control diet flies within a 5 kb window at the sweet gustatory receptor *Gr5a* and the bitter gustatory receptor *Gr66a*. **(C)** Fold change of the proportion of genes allocated to the five chromatin states (assigned in color) according to their transcription start site (TSS), normalized to the expected proportion across the whole genome (see Methods). **(D)** Mean log_2_(*Dam::Pcl/Dam*) chromatin binding coverage for a 10 kb window centered on predicted PREs on a control (gray) and sugar (purple) diet. **(E-F)** Mean CATaDa chromatin coverage normalized to 1x dm6 genome for a 10 kb window centered on **(E)** *Pcl* peaks and **(F)** predicted PREs on a control (gray) and sugar (yellow) diet. **(G)** Box plots represent the median and variance of normalized *Dam::Pcl/Dam* reads for genes differentially bound by *Pcl* on a control (gray) and sugar (purple) diet. Genes are placed into groups with higher (Group 1) or lower (Group 2) *Pcl* chromatin binding in the sugar diet environment. **(H)** GO terms associated with differentially bound genes by *Dam::Pcl/Dam* identified by iPAGE, text boxes represent the GO term category, metabolism (orange) and regulatory (lavender) (for the full iPAGE list see Supplemental Figure 4). For all panels peak FDR<0.01 (see Methods).

Most of the variation in the biological replicates of *Dam::Pcl* normalized to *Dam* alone (see Methods) was due to diet (Fig. S3A), consistent with high Pearson correlations within each dietary condition (Fig. S3B). Further, the chromatin accessibility profile of *Dam* at the *Gr5a* sweet taste receptor gene promoter was high, while at the *Gr66a* bitter taste receptor promoter –which is only expressed in bitter cells, closely located near the sweet cells– was low (Fig. 3B), suggesting that the transgenes were appropriately targeted to the sweet taste neurons and that the limited induction controlled for background DNA methylation.

We first analyzed *Pcl* chromatin occupancy in the *Gr5a+* neurons of flies on a control diet by comparing our data to a previous study that annotated five major chromatin types in *D. melanogaster* using a similar technique (DNA Adenine Methyltransferase Identification, Dam-ID) (*30*). *Pcl* targets were enriched in Polycomb chromatin (blue), compared to other chromatin types (green and black = repressive; red and yellow= active) (Fig. 3C); for example, *Pcl* occupancy was high and *Dam* accessibility low at two known Polycomb blue chromatin clusters, (Fig. S3E), while the opposite was true for regions in other chromatin types (Fig. S3F, red and yellow) (*30*). We next asked whether *Pcl* was enriched at Polycomb Response Elements (PREs), cis-regulatory sequences to which Polycomb Group Proteins bind in *D. melanogaster* (*31, 32*). Using a recently developed tool that predicts PRE regions genome-wide (*33*), we found that *Pcl* was present in these regions (Fig. 3D, gray line), with an enrichment for intergenic (3.2-fold enriched, *p*<0.001, Monte Carlo permutation test) and enhancer PREs (2.9-fold enriched, *p*<0.001, Monte Carlo permutation test).

To determine changes in *Pcl* occupancy with diet, we compared *Pcl* chromatin binding between flies fed a control and sugar diet. While ∼70% of the overall *Pcl* peaks were shared between the control (CD) and sugar diet (SD) (Fig. S3C, D), we found more *Pcl* at PREs on a sugar diet compared to a control diet (Fig. 3D, purple line). Interestingly, chromatin accessibility at both *Pcl* peaks (Fig. 3E) and PREs (Fig. 3F) was decreased in the sugar diet condition. Our analysis also showed that differentially bound *Pcl* peaks had a 3.3-fold enrichment of overlap for enhancer-type PREs (p<0.001, Monte Carlo permutation test). Further examination of the differentially bound genes, revealed a redistribution in *Pcl* occupancy (Fig. S3G), with a similar number of genes with higher (group 1) and lower (group 2) *Pcl* binding on a sugar diet compared to the control diet (Fig. 3G). Using iPAGE for pathway enrichment analysis (*34*) we found that most of the genes differentially bound by *Pcl* were transcription factors with both promoter and enhancer binding. Notably, transcription factors implicated in axon target recognition and nervous system development showed an enrichment in *Pcl* binding on a sugar diet, while those involved in proboscis development and feeding behavior had both an increase and decrease in *Pcl* occupancy on a sugar diet (Fig. 3H, for full iPAGE analysis see Fig. S4). While the large majority of genes differentially bound by *Pcl* were in the gene regulation category (80%), the pathway enrichment analysis also uncovered a few metabolism Gene Ontology (GO) terms (Fig. 3H and Fig. S4). In summary, we found that PRC2.1 targeted transcription factors involved in neuronal processes and development in the *Gr5a+* neurons, and its chromatin occupancy was redistributed at these loci in the high sugar diet environment. This redistribution could result in changes in the expression of these transcription factors and their targets, and in turn, affect the responsiveness of the sensory neurons and taste sensation.

### PRC2.1 sculpts the transcriptional responses of the *Gr5a+* neurons in response to diet

To test the hypothesis that redistribution of PRC2.1 chromatin occupancy alters the physiology of the sweet sensing neurons by changing gene expression, we used Translating mRNA Affinity Purification (TRAP) (*35*) to isolate mRNAs associated with the ribosomes of these cells. (Fig. 4A). To capture the dynamics of this process, we collected samples from age-matched *Gr5a>Rpl3-3XFLAG* flies fed a sugar diet for 3 and 7 days (Fig. S5A). We first verified that this technique selected for mRNAs in the *Gr5a+* neurons alone by quantifying the normalized read counts (*Gr5a+*/input) for three sweet taste receptor genes *Gr5a*, *Gr64f*, and *Gr64a,* which are expressed in cells labeled by the *Gr5a*-GAL4. Indeed, these transcripts were enriched in the *Gr5a+* fraction compared to the input (Fig. S5B), while the opposite was true for the bitter receptor gene *Gr66a*, which is expressed only in the bitter sensing neurons in the taste sensilla (*36*) (Fig. S5B).

**Fig. 4.**
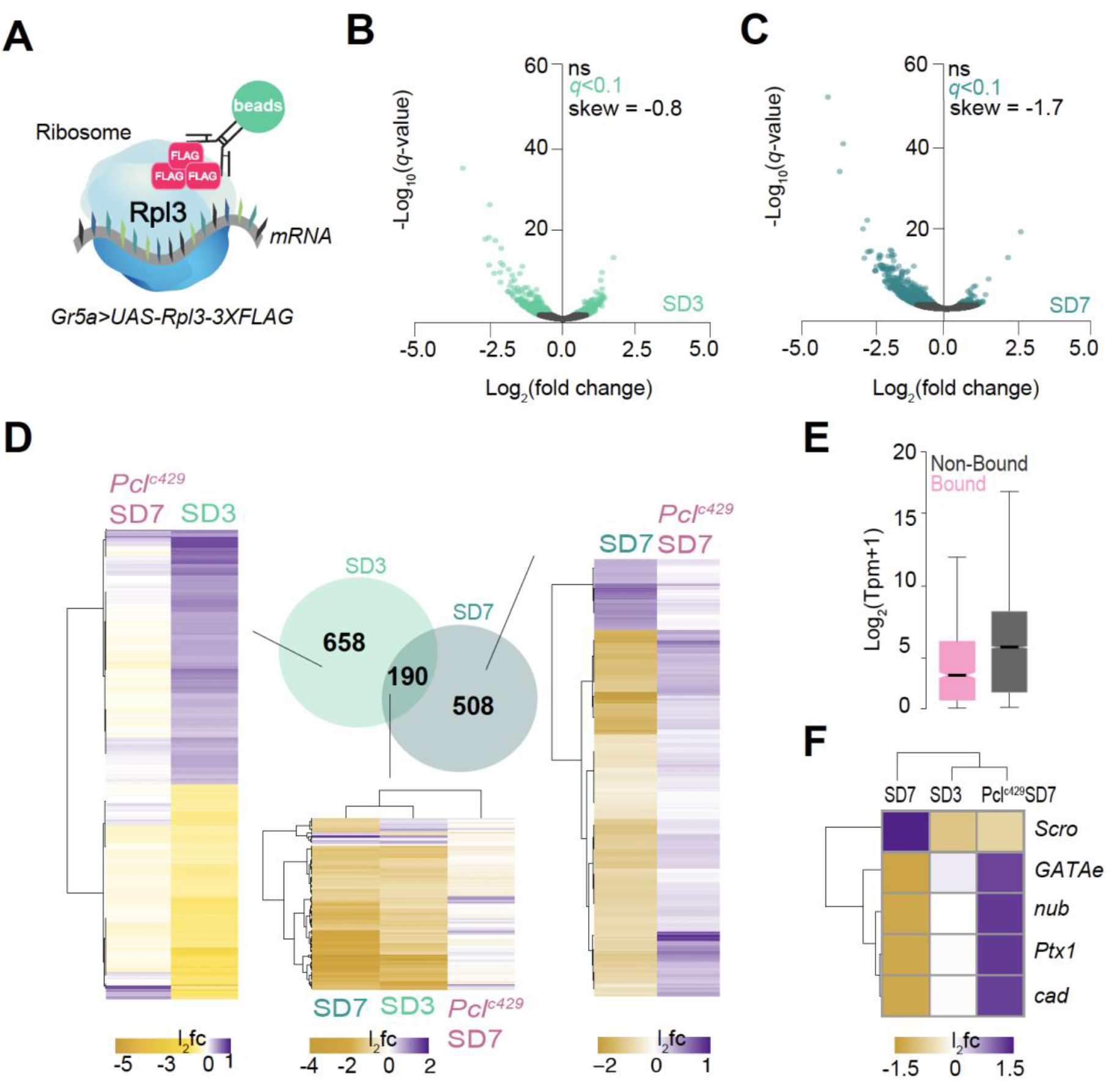
PRC2.1 sculpts the transcriptional responses of the *Gr5a+* neurons in response to diet. **(A)** Schematic depiction of the TRAP technique used to profile the changes in the *Gr5a+* neurons. **(B-C)** Volcano plots representing changes in gene expression in the *Gr5a+* neurons of age matched male *Gr5a>UAS-Rpl3-3XFLAG* flies on a sugar diet for 3 days (SD3, mint) and 7 days (SD7, teal). n=2-3 replicates per condition (10,000 *Gr5a+* cells per replicate). Non-significant genes are shown in black and genes with *q*<0.1 (Wald test) are in mint or teal for SD3 and SD7 respectively, comparison is to the control diet group. **(D)** Venn diagram of differentially expressed genes (DEGs) at SD3/CD (mint), SD7/CD (teal), and the overlap between SD3/SD7 (Wald test, *q*<0.1). Heatmaps show log_2_ fold changes (l_2_fc) for DEGs in each condition specified in Venn diagram (left columns in heatmap, SD3, SD7, and SD3+SD7, all comparisons to control diet) and *Pcl_c429_* mutant flies (right column in heatmap, *Pcl_c429_ SD7* comparison to control diet); l_2_fc range from purple (high) to gold (low). **(E)** Normalized read counts (Tpm+1) from TRAP for *Pcl* bound (pink) and not bound genes (gray) on a control diet. Box plots represent the median and variance, *p*-value = 3.196×10_-6_, two-tailed t-test. **(F)** Log_2_ fold changes for *scarecrow* (*Scro*), *Paired-like homeobox domain 1* (*Ptx1*), *caudal* (*cad*), *GATAe,* and *nubbin* (*nub*) in control SD7/CD, SD3/CD flies, and *Pcl_c429_*SD7/CD mutant flies. L_2_fc ranges from purple (high) to gold (low).

Notably, we observed a large negative skew in gene expression in the *Gr5a*+ neurons of flies fed a sugar diet for 3 (SD3, mint; compared to the control diet) and 7 days (SD7, teal; compared to the control diet) (Fig. 4B-C, −0.8 and −1.7 skew), with ∼68% and 87% of differentially expressed genes (DEG, each compared to control diet, Wald test, *q*< 0.1) showing negative log_2_ fold changes (l_2_fc) respectively (Fig. 4B-C, SD3 and SD7), consistent with the idea that a repressive gene regulatory mechanism is involved in this process. Overall, we found ∼800 transcripts differentially expressed at each time point compared to the control diet, while ∼190 changed at both time points (Fig. 4D, Venn diagram, Wald test, *q*< 0.1). Gene Ontology (GO) analysis (*34*) revealed that these genes were part of biological pathways involved in 3 broad categories: neural function/signaling, metabolism, and gene regulation (Fig. S6 and Fig. S7). GO terms for neuron-specific processes, such as dendritic membrane, sensory perception of chemical stimulus, and presynaptic/vesicle transport, were enriched at both timepoints (Fig. S6 and Fig. S7), suggesting that PRC2.1 may alter the physiology of the sensory neurons through these pathways in response to a high sugar environment. Interestingly, flies fed a sugar diet for 7 days also showed changes in GO terms linked to neurodevelopmental processes, such as asymmetric neuroblast division and neuron projection morphogenesis, indicating that longer exposure to the diet led to additional alterations in neural function, which may explain the worsening of sweet taste sensation at the later time point (*18*). GO terms associated with metabolic changes were present in higher numbers in flies fed a sugar diet for 7 days, consistent with the findings that longer exposure to the high sugar diet leads to higher fat accumulation (*18*). Finally, we observed changes in “regulatory” GO terms such as transcription factor and corepressor, consistent with the redistribution of *Pcl* chromatin occupancy in response to a high sugar diet. Thus, consumption of a high sugar diet altered neural, regulatory, and metabolic genes in the *Gr5a+* cells.

To determine the role of PRC2.1 in these changes, we performed the transcriptional profiling experiments in the *Gr5a+* neurons of *Pcl_c429_* mutant animals fed a control diet and sugar diet for 7 days (CD and SD7) (Fig. S5C). Strikingly, the *Pcl* mutation abolished the negative skew (Fig. S5D) and largely nullified the effects of the high sugar diet environment on gene expression. Specifically, of the genes repressed by a sugar diet (Fig. 4D, heatmap) 32% had a positive log_2_fold change (Wald test, *q*< 0.1) and 76% were unchanged (*q*<0.1, practical equivalence test using a null hypothesis of a change of at least 1.5-fold; see Methods for details) between *Pcl* mutants fed a control and sugar diet. This effect was reflected in the GO analysis where terms changed by a high sugar diet in *Pcl* wild-type animals, such as dendritic membrane, sensory perception of chemical stimuli, synapse, and carbohydrate metabolic process, showed opposite trends in log_2_ fold changes in *Pcl* mutants (Fig. S8). Thus, *Pcl* mutations abolished nearly all the gene expression changes induced by a high sugar diet consistent with their effects on behavior (PER, Fig.1), neural function (*in vivo* calcium imaging, Fig. 2), and metabolism (triglycerides, Fig. 2). Together, these findings support the hypothesis that PRC2.1 tunes taste sensation to the food environment by influencing the expression of genes involved in the physiology of the sensory neurons.

### PRC2.1 represses a transcriptional program required for sweet taste

We discovered that a high sugar diet environment repressed gene expression in the sweet sensory neurons, and that *Pcl* mutations almost entirely abolished this effect. This, together with the discovery that *Pcl* binding primarily changed at the enhancers of transcription factor genes on a sugar diet, suggests that *Pcl* redistribution may affect the expression of transcription factors that, in turn, control genes responsible for the overall responsiveness of these sensory neurons to sweetness. This idea was supported by the observation that *Pcl*-bound genes had lower expression levels than those not bound by it in the *Gr5a+* neurons (Fig. 4E), with many genes showing high binding and low expression (log_2_tpm <2, dark purple), while others having higher mRNA read counts (log_2_tpm >5, light purple) (Fig. S5E). To test this hypothesis, we looked for transcription factors that were differentially bound by *Pcl* directly and that showed changes in gene expression on a sugar diet (Fig. S5F). This analysis revealed 5 transcription factors: four of these were activators – *GATAe* (Zn finger)*, nubbin/pdm* (*nub*, POU homeobox)*, Ptx1* (paired-domain homeobox), and *caudal* (*cad*, hox-like homeobox)– which had higher *Pcl* binding (Fig. 5A) and lower mRNA levels on a sugar diet (Fig. 4F). The fifth transcription factor was the suppressor *scarecrow* (*Scro*, NK-like homeobox), which had lower *Pcl* binding (Fig. 5A) and higher mRNAs levels on a sugar diet (Fig. 4F). Notably, mutations in *Pcl* reversed the effects of a high sugar diet on the expression of these 5 genes, suggesting that the binding of *Pcl* modulates their mRNA levels (Fig. 4F). Interestingly, *cad, Ptx1, GATAe,* and *nub* were enriched in the *Gr5a+* neurons compared to the input (heads), while *Scro* was depleted in these cells (Fig. S9A). To test the effects of these five transcription factors on sweet taste, we manipulated their expression to mimic the direction of change on a high sugar diet. Knockdown of *cad, Ptx1, GATAe,* or *nub*, and overexpression of *Scro* in the *Gr5a+* neurons of flies fed a control diet led to a decrease in sweet taste sensation (Fig. 5B) comparable to that experienced by animals on a sugar diet (Fig. 1B). Thus, *cad, Ptx1, GATAe,* and *nub* are direct targets of PRC2.1 and necessary for sweet taste, while overexpression of *Scro* is sufficient to decrease it. However, overexpression of each activator alone and knockdown of *Scro* was not sufficient to rescue sweet taste in flies fed a sugar diet (Fig. S9B).

**Fig. 5.**
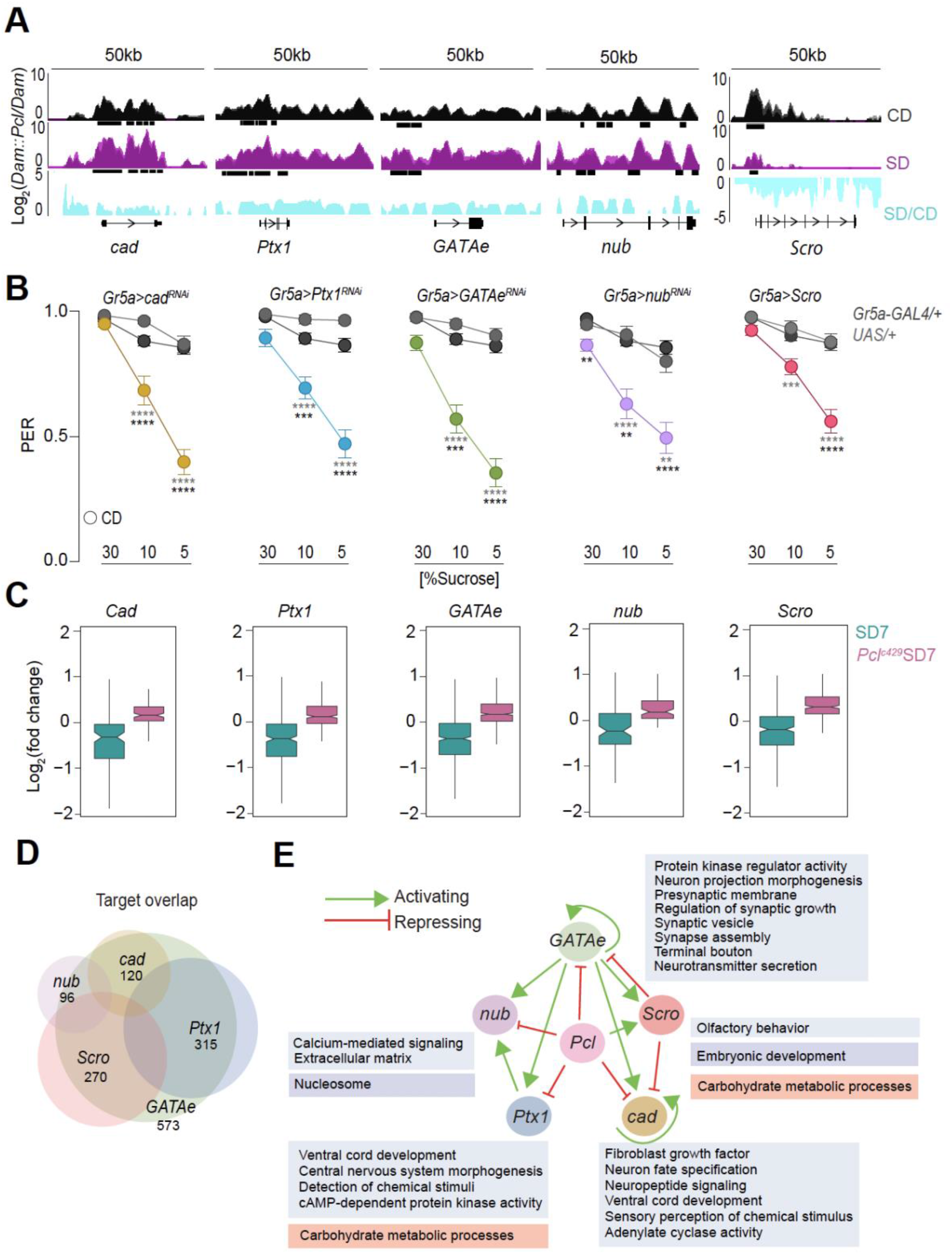
PRC2.1 represses a transcriptional program required for sweet taste. **(A)** *Log_2_(Dam::Pcl/Dam*) chromatin binding coverage tracks of flies on a control (black) and sugar (purple) diet within a 50 kb window at the *cad*, *Ptx1*, *GATAe*, *nub*, and *Scro* genes visualized with the UCSD genome browser. The three biological replicates for each condition are superimposed. Turquoise traces are SD/CD fold changes. Peaks are shown as black boxes below respective traces (*q*<0.01) and genes are shown in dense format to include all potential isoforms. **(B)** Taste responses (y axis) to stimulation of the labellum with 30%, 10% and 5% sucrose (x axis) in age-matched males of *Gr5a>cad_RNAi_* (gold), *Gr5a>Ptx1_RNAi_* (blue), *Gr5a>GATAe_RNAi_* (green), *Gr5a>nub_RNAi_* (lavender), *Gr5a>Scro* (salmon), and parental transgenic control (gray, crossed to *w1118_cs_*) flies on a control diet for 7 days. n = 30-54, Kruskal-Wallis Dunn’s multiple comparisons, comparisons to both transgenic controls. All data shown as mean ± SEM, ∗∗∗∗*p* < 0.0001, ∗∗∗*p* < 0.001, ∗∗*p* < 0.01, and ∗*p* < 0.05. **(C)** Box plots represent the median and variance of the log_2_ fold changes (l_2_fc) for candidate gene targets of *cad*, *Ptx1*, *GATAe*, *nub*, and *Scro* (for threshold cutoff score see Supplemental Figure 9) in control flies at SD7 (teal), and *Pcl_c429_* mutants at SD7 (pink). **(D)** Venn diagram representation of the overlap of the candidate gene targets of *cad* (gold), *Ptx1* (blue), *GATAe* (green), *nub* (lavender), and *Scro* (salmon). **(E)** A transcriptional loop between *cad* (gold), *Ptx1* (blue), *GATAe* (green), *nub* (lavender), and *Scro* (salmon) mediated by *Pcl.* Green arrows represent activation and red bars represent inhibition. GO terms associated with each transcriptional regulator identified by iPAGE are listed. Text boxes represent the GO term category, metabolism (orange), regulatory (lavender) and neural/signal (blue) (for the full iPAGE list see Supplemental File 5).

Given that the 4 activators are required for sweet taste sensation, we reasoned that they may control the expression of genes important for the proper function of the *Gr5a+* neurons and normal sweet taste. To identify candidate target genes, we tiled the entire *D. melanogaster* genome using the motifs for each of these 5 transcription factors, converted the hits for each transcription factor to z-scores, and determined candidate regulatory targets based on estimates of the z-score threshold for binding in each case (Fig. S9C; see Methods for details). We then flagged as “targets” the set of genes that had a putative binding site (exceeding our transcription factors-specific z-score cutoff) within a 2 kb region upstream of the annotated ORF start (Fig. S9D-H), and examined their expression pattern in the *Gr5a+* neurons of flies on a control and sugar diet. This analysis revealed 658 genes that were collectively regulated by these five transcription factors. Targets of the transcriptional activators Cad, Ptx1, GATAe and Nub– which *Pcl* repressed on a sugar diet– showed negative log_2_fold changes on a sugar diet (Fig. 5C, SD7, teal) that reverted in the *Pcl* mutants (Fig. 5C, pink). Conversely, targets of the transcriptional repressor Scro – which was released from *Pcl* binding and had higher mRNA levels on a sugar diet – showed negative log_2_fold changes on a sugar diet (Fig. 5C, SD7, teal) that reverted in *Pcl* mutants (Fig. 5C, pink). Strikingly, the 658 putative targets of these 5 transcription factors accounted for nearly all the genes changed by a high sugar diet (Fig. 4), suggesting that by directly modulating the expression of *Ptx1, cad, GATAe, nub,* and *Scro*, PRC2.1 could influence the neural state of the *Gr5a+* neurons via their targets. When we examined whether these targets were regulated by more than one of the 5 transcription factors targeted by *Pcl*, we found significant overlap among the regulons of all of the 4 transcriptional activators with the exception of *Ptx1-nub* (Fisher’s exact test, FDR-corrected *p*<0.000001) (Fig. 5D), suggesting that they could direct the expression of a common set of genes. Further, a portion of *Scro* targets also overlapped with the regulons of each of the activators (Fisher’s exact test, *q*<0.000001), indicating that *Scro* could further drive the negative skew in gene expression on a high sugar diet beyond that caused by the direct binding of PRC2.1 to the activators. Transcription factors that share common targets are often part of feed-forward loops, where they regulate one another and themselves to ensure stability of gene expression. Indeed, we found that *GATAe* had binding sites in the promoters of all four regulators considered here (*cad*, *Scro*, *Ptx1*, *nub*), in addition to binding its own promoter in an auto-regulatory loop (Fig. 5E). Furthermore, Cad also targeted itself, *nub* was one of Ptx1 targets, and Scro regulated both *cad* and *GATAe*, forming a negative feedback loop with the latter (Fig. 5D). Thus, the 5 transcription factors differentially bound by PRC2.1 on a high sugar diet, form a hub that regulates the physiology of the *Gr5a+* neurons. To gain a deeper understanding of which aspects of physiology were changed, we used pathway enrichment analysis on the regulons for each transcription factor. GATAe targets, which comprise a large number of the genes regulated by the 4 other transcription factors, were enriched for GO terms involved in synaptic assembly and growth, terminal bouton, neural projection morphogenesis, and protein kinase regulation (summarized in Fig. 5D, and Supplementary File 5). In contrast, Ptx1 targets were enriched in GO terms implicated in cyclic AMP signaling, detection of chemical stimuli, and morphogenesis (summarized in Fig. 5D, and Supplementary File 5), Cad targets showed enrichments in adenylate cyclase activity, sensory perception, and neuropeptide signaling (summarized in Fig. 5D, and Supplementary File 5), and Nub targets in calcium signaling and nucleosome (summarized in Fig. 5D, and Supplementary File 5). The targets of the repressor Scro showed enrichment in both neural and metabolic GO terms such as olfactory behavior and carbohydrate metabolic process (summarized in Fig. 5D, and Supplementary File 5).

To test the possibility that these targets form a functional network, we used STRING (*37*) and found a significant number of interactions above the expected number (protein-protein interaction enrichment of *p* < 1.0e-16) suggesting that the targets are indeed part of a functional and biologically connected network in the *Gr5a+* neurons. We then used a subset of the neural targets to build a second network to identify candidate target genes that may play a direct role in neural physiology and sweet taste. This network showed strong interactions between genes involved in synaptic organization and signal transduction and their connection with the upstream regulators (Fig. S14A). We chose two genes at the edges of the network, which are less likely to have redundant functions, the *Adenylyl Cyclase X D* (*ACXD*) gene (*38*) and the *Activity Regulated Cytoskeleton Associated Protein 1* (*Arc1*) (*39*), which are involved in the sensing of stimuli and in synaptic plasticity, respectively. Knockdown and mutations of *Arc1* or *ACXD* in the *Gr5a+* cells of flies on a control diet led to a significant decrease in sweet taste responses compared to the transgenic controls (Fig. S14B-D). Together, these lines of evidence suggest that PRC2.1 mediates the effects of a high sugar diet on sweet taste by directly controlling the expression of a transcription program required for sweet taste.

### The persistent phenotypic memory of the food environment is dependent on PRC2.1

The cellular fates created by Polycomb Group proteins are inherited as stable memories across cell divisions to ensure phenotypic stability even in the absence of the triggering stimuli (*40–44*). We therefore asked if the neural and behavioral state created by PRC2.1 in the high sugar diet environment was maintained when files were moved to the control diet after eating a sugar diet for 7 days (SD>CD) (Fig. 6A). We found that these animals had lower sweet taste, similar to that of age-matched flies fed a sugar diet for 7 days (Fig.6B, SD>CD compared to CD>SD). However, their triglyceride levels were similar to those of control diet flies (Fig. S14E), suggesting that while fat storage was reversible when flies only had access to the “healthy” control diet, sweet taste deficits persisted.

**Fig. 6.**
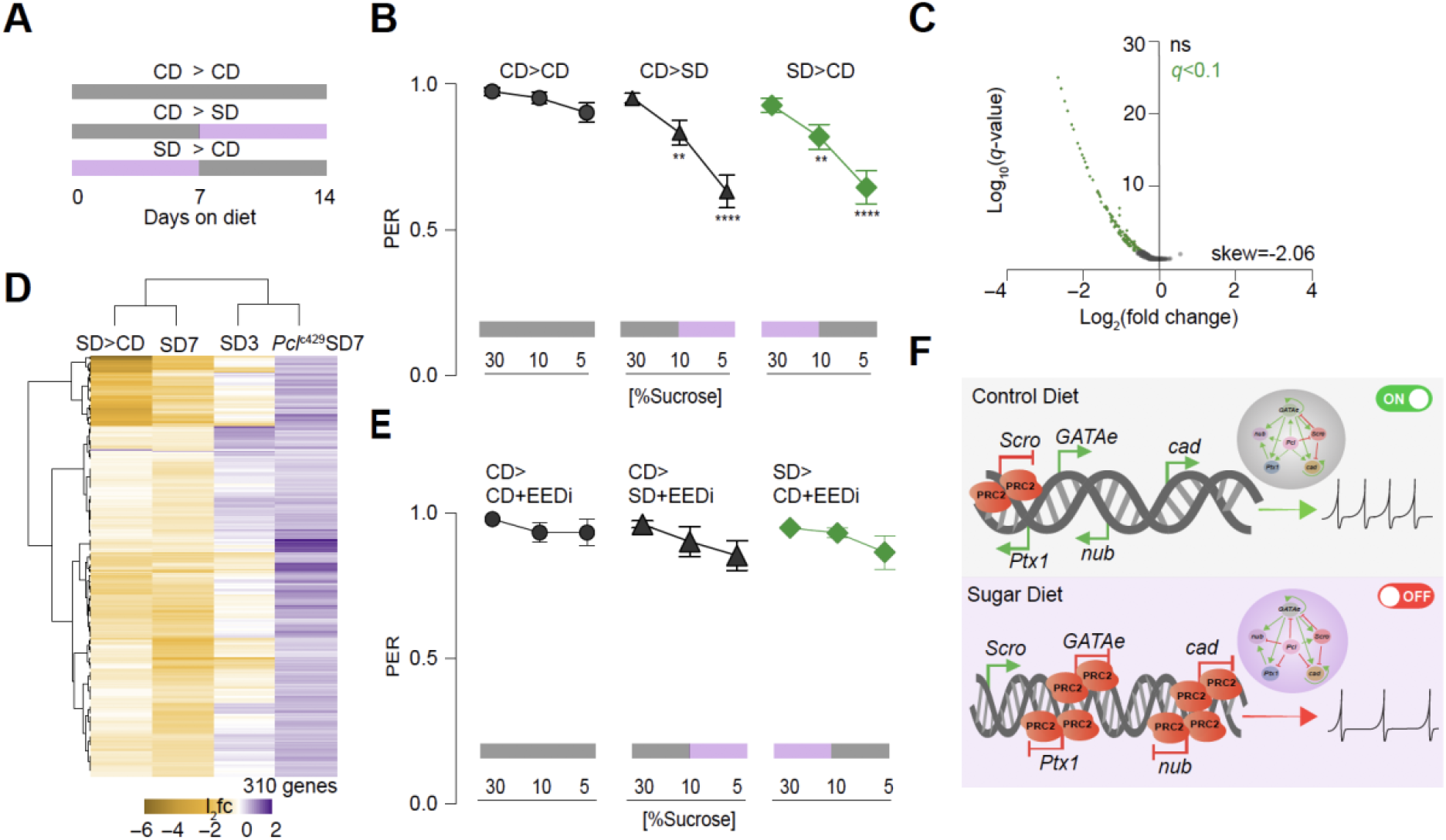
The persistent phenotypic memory of the food environment is dependent on PRC2.1. **(A)** Diagram representing dietary manipulations, CD, control diet (gray), SD, sugar diet (purple), and > represents the switch in diet. **(B)** Taste responses (y axis) to stimulation of the labellum with 30%, 10% and 5% sucrose (x axis) of age-matched males *w1118_cs_* flies on a control (CD>CD, circle), control to sugar (CD>SD, triangle), and sugar to control (SD>CD, diamond) diet as shown in (A) n =57-64, Kruskal-Wallis Dunn’s multiple comparisons, comparisons to control diet. All data shown as mean ± SEM, ∗∗∗∗*p* < 0.0001, ∗∗∗*p* < 0.001, ∗∗*p* < 0.01, and ∗*p* < 0.05 in this panel. **(C)** Volcano plot with the changes in gene expression in the *Gr5a+* neurons of age-matched male *Gr5a>UAS-Rpl3-3XFLAG* flies on a sugar to control (SD>CD, green) diet compared to the control diet group (CD>CD, Wald test, *q*<0.1, ns is non-significant). n=2 replicates per condition (∼10,000 *Gr5a+* cells per replicate). **(D)** Heatmap showing how the differentially expressed genes identified in Figure 5C are changing in the SD>CD condition (310/658, 48% Wald test, *q*<0.1) compared to the other conditions SD3, SD7, and *Pcl_c429_*SD7 (column headings, all comparisons to control diet). Log_2_fold changes for these genes were determined by comparing each group to its own control (see Methods). L_2_fc ranges from purple (high) to gold (low). **(E)** Taste responses (y axis) to stimulation of the labellum with 30%, 10% and 5% sucrose (x axis) of age-matched males *w1118_cs_* flies on a control (circle), control to sugar (triangle), and sugar to control (diamond, green) diet for 14 days. EEDi was supplemented to the diets at the day 7 transition (see diagram in figure). n =34-46, Kruskal-Wallis Dunn’s multiple comparisons, comparisons to control diet. All data shown as mean ± SEM, ∗∗∗∗*p* < 0.0001, ∗∗∗*p* < 0.001, ∗∗*p* < 0.01, and ∗*p* < 0.05 in this panel. **(F)** Model of molecular changes in the *Gr5a+* neurons on a control (top panel) and sugar (bottom panel) diet, showing the redistribution in PRC2.1 binding, and the effects on the transcription of the regulators (green and red arrows), and neural responses to sweet stimuli.

To understand how this phenotype compares to that of the control diet flies at the molecular level, we conducted TRAP of the *Gr5a+* neurons of flies in the SD>CD and CD>CD conditions. mRNAs from flies on a SD>CD showed a strong negative skew in log_2_fold changes compared to the control diet group (Fig. 6C, −2.02), reminiscent of the skew we observed in flies fed a sugar diet (Fig. 4C). Furthermore, we observed that 47% (310/658) of genes in the transcriptional network repressed by PRC2.1 on a sugar diet were still decreased in SD>CD flies (Fig. 6D). Interestingly, the SD>CD animals clustered with the sugar diet 7 (SD7) group compared to sugar diet 3 (SD3) and *Pcl* mutants fed a sugar diet (Fig. 6D). Thus, at the molecular level, half of the neural state established by dietary sugar via PRC2.1 persisted. To test the hypothesis that PRC2.1 plays an active role in the maintenance of this neural state in the absence of the sugar diet, we inhibited PRC2 activity during the “recovery” diet using an EEDi inhibitor (SD>CD+EEDi). Remarkably, these animals showed a restoration of wild-type sweet taste (Fig. 6E, green triangles) compared to flies fed the recovery diet without EEDi supplementation (Fig. 6B, SD>CD). Together, these data indicate that the sensory neurons retain a transcriptional and phenotypic memory of the sugar diet environment that leads to long lasting behavioral deficits, and that PRC2.1 is constitutively required for the persistence of this state.

## Discussion

In this study we set out to understand how diet composition alters the gustatory system to promote food intake and weight gain. Specifically, we took advantage of the simple sensory system of *D. melanogaster* and its exquisite genetic and neural tools to identify the molecular mechanisms through which diet composition changes neural state, physiology, and behavior. Here we show that the decrease in sweet taste sensation that flies experience after chronic exposure to a high sugar diet is caused by the cell-autonomous action of the Polycomb Repressive Complex 2.1 in the sweet gustatory neurons. Mutations and pharmacological inhibition of PRC2.1 blocked the effects of the food environment on neural activity, behavior, and obesity. While we do not exclude the possibility that PRC1 and PhoRC may also be involved, we found that mutations or knockdown in these complexes had no effect on taste. In the high sugar food environment, PRC2.1 chromatin occupancy was redistributed, leading to the repression of transcription factors, neural, signaling, and metabolic genes that decreased the responsiveness of the *Gr5a+* neurons and the fly’s sensory experience. However, we discovered that PRC2.1 did not directly bind to neuronal genes in these cells and that, instead, it targeted transcription factors involved in sensory neuron development, synaptic function, and axon targeting. Specifically, on a high sugar diet *Pcl* binding was increased at the *cad*, *GATAe*, *nub/pdm*, *Ptx1* loci and decreased at the *Scro* locus, with corresponding changes in the mRNA levels of these genes (Fig. 6F, model). Together, the decrease in the mRNA levels of the 4 activators (*cad*, *GATAe*, *nub/pdm*, *Ptx1*) and the increase in the repressor (*Scro*), led to the repression of genes implicated in synaptic function, signal transduction, and metabolism. Interestingly, we uncovered several feed-forward regulatory loops among the transcriptional regulators, suggesting that this hub could ensure the stability of the neural state unless the change in environment is chronic. This transcriptional network controlled by these regulators was necessary for sweet taste, because knockdown of the four activators and a few of their targets, as well as overexpression of *Scro* (Fig. 5B and Fig. S10B-D), resulted in a decrease in sweet taste on a control diet.

Interestingly, several of the transcription factors we identified – *Ptx1*, *Scro*, and *nub/pdm* – have been shown to control the proper branching, synaptic connectivity, and function of sensory neurons (*45–51*), while others (*cad*, *nub/pdm*) play a role in neuroblast development (*52, 53*); PRC2 also functions as a competence factor in neural proliferation, differentiation and sensory neurons (*45, 52, 54*). Importantly, we found that the 4 activators that are repressed by *Pcl* in the high sugar condition are enriched in the *Gr5a+* cells on a control diet, while *Scro* is depleted. Thus, the transcriptional network of ∼658 genes controlled by this developmental transcription hub is likely to define the intrinsic properties of the sweet sensing neurons. Indeed, we observed that many of the target genes involved in signaling, synaptic function, and cell adhesion such as the kinase *haspin*, the adenylate cyclase *ACXD*, *sytalpha*, *Arc1*, tetraspanin, jonan, and innexin proteins among others, were part of highly interconnected network, which we speculate, could affect the sweet gustatory neurons at both the functional and connectivity levels. Since we did not observe a change in the expression of the sweet taste receptors, or the misexpression of other taste receptors (*36*), our data are not consistent with a complete loss of identity of the *Gr5a+* neurons. Instead, we propose that PRC2.1 tunes the sweet sensory neurons to the dietary environment by altering a transcriptional network that controls the intrinsic properties of these cells, especially those involved in signal transduction, connectivity, synaptic function, and metabolism. Studies that test the effects of this network on the connectivity, morphology, and signal transduction of the sweet sensory neurons will shed light on how exactly the transcriptional remodeling caused by PRC2.1 impacts the *Gr5a+* cells. The question of how exactly PRC2.1 senses the changes in the food environment also remains open. Recent studies suggest that the activity of Polycomb Group proteins is directly and indirectly linked to cellular metabolism, including kinase signaling cascades, GlcNAcylation, and the availability of cofactors for histone modifications (*55–59*). Our experiments show that PRC2.1 chromatin occupancy changes depending on the dietary environment, and provides a new model to tackle this question. Finally, dysregulation of Polycomb-associated chromatin has been found in mice and humans with diet-induced obesity (*60–62*), suggesting that the mechanisms we discovered here could also underlie the chemosensory alterations reported in mammals.

More broadly our work opens up the exciting possibility that PRC2 may modulate neural plasticity in response to environmental conditions. Despite its central role in development and maintenance of neural identity, studies have not directly linked PRC2.1 to neural plasticity. However, in other post-mitotic cells such as muscle, Polycomb Group Proteins are known to reshape transcriptional programs according to environmental stressors, such as oxidative stress, injury, temperature, and light (*55, 56*). Our findings advance the conceptual understanding of the role of Polycomb Group Proteins in the nervous system and argue that PRC2 could regulate “neural states” and metaplasticity in response to environment stimuli beyond diet. Using neuroepigenetic mechanisms like those employed by Polycomb Group Proteins to tune neural states to external conditions could provide several advantages compared to the medley of other cellular, receptor, or synaptic plasticity based mechanisms. Specifically, it would allow cells to 1) orchestrate a coordinated response, 2) create a memory of the environment, and 3) buffer small fluctuations until a substantial challenge is perceived. It is particularly fascinating to think about the molecular mechanisms through which these neural states may be established. The need of neurons to constantly maintain their identity may mean that environmental signals like the extent of sensory stimulation could alter the expression of developmental gene batteries and affect neural states (*63*). Indeed, it has been speculated that some forms of plasticity may re-engage developmental programs that specify the intrinsic properties of neurons (*64, 65*). Here we observed that the regulators of the transcriptional network we uncovered function in sensory neuron development and are enriched in the *Gr5a+* cells. Thus, it could be a hallmark of neuroepigenetic plasticity to exploit developmental programs, linking the known role of PRC2 in establishing cell fates with this newly discovered function in modulating cell states. Incidentally, engaging developmental programs could be the reason why some environments and experiences leave a memory that leads to the persistent expression of the phenotype beyond the presence of the triggering stimulus, as these could target neural connectivity and set synaptic weights thresholds. Defining the circuit-specific changes of the pioneering studies on maternal care, addiction, and learning and memory (*66–69*) would put this hypothesis to test. Here we found that the changes in taste sensation and half of the sugar diet neural state set by PRC2.1 remained even after animals were moved back to the control diet for a week. A limitation of our study is that due to the small number of *Gr5a+* neurons and their anatomically inaccessible location, we were not able to measure the identity of the molecular memory in these cells alone. However, we saw that the phenotypic memory of the high sugar food environment was dependent on the constitutive action of PRC2.1. Based on other studies showing that the H3K27 methyl mark acts as a molecular memory during development (*40, 41*), we speculate that this is likely to be the molecular signal in the *Gr5a+* cells too.

In conclusion, we show that PRC2.1 mediates the effects of dietary sugar on sweet taste by establishing persistent alterations in the taste neurons that remain as a phenotypic and neural memory of the previous food environment. We speculate that this memory may lock animals into patterns of feeding behavior that become maladaptive and promote obesity. Thus, diet can induce lasting molecular alterations that restrict the behavioral plasticity of animals and impact disease risk. Since the content of sugar in processed foods is similar to that we fed flies and the function of Polycomb Group Proteins is conserved from plants to humans (*20*), our work is broadly relevant to understanding the effects of processed food on the mammalian taste system and its impact on food intake and a whole range of diet-related conditions and diseases that affect billions of people around the globe.

## Materials and Methods

### Fly Husbandry and Strains

All flies were grown and maintained on cornmeal food (Bloomington Food B recipe) at 25°C and 45%– 55% humidity under a 12:12 hour light-dark cycle (ZT0 at 9 AM). Male flies were collected under CO2 anesthesia 1-3 days after eclosion and maintained in a vial that housed 35-40 flies. Flies were acclimated to their new vial environment for an additional 2 days. For all experiments, flies were changed to fresh food vials every other day.

For all dietary manipulations, the following compounds were mixed into standard cornmeal food (Bloomington Food B recipe) (0.58 calories per gram) by melting, mixing, and pouring new vials as in (*70*) and (*71*). For the 30% sugar diet (1.41 calories per gram) Domino granulated sugar w/v was added. For the EED226 inhibitor diet (AxonMedchem), EED226 was solubilized in 10% DMSO and added to control or 30% sugar diet at a total concentration of 8 uM.

For genetic manipulations the GAL4/UAS system was used to express transgenes of interest in *Gustatory receptor 5a Gr5a-GAL4*. For each GAL4/UAS cross, transgenic controls were made by crossing the *w1118_CS_* (gift from A. Simon) to GAL4 or UAS flies, sex-matched to those used in the GAL4/UAS cross. The following fly lines are used in this paper: *Gr5a-GAL4/Cyo* (gift from K. Scott, University of California, Berkeley), *nsyb-GAL4/Cyo* (gift from J. Simpson, University of California, Santa Barbara), *Gr64f-GAL4/Cyo* (gift from H. Amrein, University of Texas, A&M), *UAS-GCaMP6s-Brp-mCherry* (BDSC #77131), *Tubulin-GAL80_ts_* (BDSC #7018), *UAS-Ogt-RNAi* and *UAS-Ogt-GFP/Cyo* (gift from C. Lehner, University of Zurich), *UAS-Pcl* (FlyORF #F001897), *UAS-Pcl-RNAi* (VDRC #v220046, BDSC #33945), *UAS-Rpl3-3XFLAG* (gift from D. Dickman, University of Southern California), *Pcl_T1_/CKG* (gift from G. Cavalli, Université de Montpellier), *Pcl_c241_/Cyo*, *E(z)_c249_/TM3*, *Su(z)12_c253_*, *Esc_c289_* (gift from N. Liu, Chinese Academy of Sciences), *Pc_1_/TM1* (BDSC #1728), *ph-d _401_ph-p_602_/FM7* (BDSC #5444), *Psc_h27_/Cyo* (BDSC #5547), *UAS-LT3-Dam* (gift from A.H Brand, University of Cambridge), *UAS-LT3-Dam::PCL* (made in this study), *UAS-Arc1-RNAi* (BDSC #25954), *Arc_esm13_* (BDSC #37530), *Arc1_esm18_*(BDSC #37531), *UAS-Arc1* (BDSC #37532), *UAS-Ptx1-RNAi* (VDRC #107785), *UAS-cad-RNAi* (BDSC #34702), *UAS-GATAe-RNAi* (BDSC #33748), *UAS-nub* (FlyORF #F000147), *UAS-Scro-RNAi* (BDSC #29387), *UAS-Ptx1* (FlyORF #F003469), *UAS-Scro* (FlyORF F003427), *UAS-nub-RNAi* (BDSC #55305), *UAS-cad* (FlyORF #F000471) *UAS-Jarid2-RNAi* (BDSC #32891, #26184), *UAS-ACXD-RNAi* (BDSC #62871, #35589), *UAS-jing-RNAi* (BDSC #27084), *UAS-Sfmbt-RNAi* (BDSC #32473).

### Proboscis Extension Response

Male flies were food deprived for 18-24 hours in a vial with a Kimwipe dampened with 2 mL of milliQ-filtered deionized (milliQ DI) water. Proboscis extension response (PER) was carried out as described in (*21*).

### Proboscis Immunofluorescence

Probosces were dissected in 1xPBS and fixed in 4% PFA, mounted in FocusClear (CelExplorer) on coverslips. Cell bodies were imaged using a FV1200 Olympus confocal with a 20x objective. Cells were counted using Imaris Image analysis software.

### Triglyceride Measurements

Total triglycerides normalized to total protein were measured as described in (*72*). Briefly, two flies per biological replicate were homogenized in lysis buffer (140 mM NaCl, 50 mM Tris-HCl pH 7.4, 0.1% Triton-X) containing protease inhibitor (Thermo Scientific). Lysate extract was used to determine protein and triglyceride concentrations using Pierce BCA assay (Thermo Scientific, abs 562 nm) and Triglycerides LiquiColor test (Stanbio, abs 500 nm), respectively.

### Calcium Imaging

Male flies expressing *GCaMP6s-Brp-mCherry* (*25*) in the sweet sensing neurons were food deprived for 18-24 hours. The flies for live imaging were prepared similar to LeDue et al. (*73*). Briefly, flies were fixed to a custom-printed plastic slide with paraffin wax and the proboscis waxed to an extended position. Distal leg segments were removed to prevent tarsal interference with labellar stimulation. To image the SEZ, a sugarless artificial hemolymph solution filled the well surrounding the head. Subsequently, the dorsal cuticle between the eyes was removed by microdissection to expose the brain. Each fly proboscis was tested with milliQ water before stimulating with 20% sucrose dissolved in milliQ water. Stimulus (a piece of Kimwipe soaked in tastant and held with forceps) delivery to the proboscis was manual and timed to coincide with the 100th recording sample of each time series. Imaging was carried out using an upright confocal microscope (Olympus, FluoView 1200 BX61WI), a 20x water-immersion objective and laser excitation at 488 and 543 nm. Recordings were made at 4 Hz (512 x 512 pixels). Plane of interest was kept to the most ventral neuropil regions innervated by the sweet sensing neurons.

### RNA Extraction and Quantitative RT-PCR

For all RNA extractions used for qPCR, fly heads from 10-20 flies were dissected into Trizol (Ambion) and homogenized with plastic pestles. RNA was extracted by phenol chloroform (Ambion), and precipitated by isopropanol with Glycoblue Coprecipitant (Invitrogen). RNA pellet was washed as needed with 75% ethanol and subsequently eluted in nuclease free water and treated with DNase I, according to manufacturer’s instructions (Turbo DNA-free DNA removal kit, Ambion). All steps were carried out in RNase free conditions, and RNA was stored at −80C until further processing.

Complementary DNA was synthesized by Superscript III (Invitrogen) reverse transcriptase with the addition of Ribolock RNase inhibitor (Thermo Fisher Scientific). qPCR reactions were carried out using Power SYBR Green PCR master mix (Applied Biosystems) based on manufacturer’s instructions. Primers were added at a 2.5 uM concentration. All reactions were run on a 96-well plate on the StepOnePLus Real-Time PCR System (Applied Biosystems) and quantifications were made relative to the reference gene Ribosomal protein 49 (Rp49). Primer sequences are listed below and were tested for efficiency prior to the qPCR experiment:

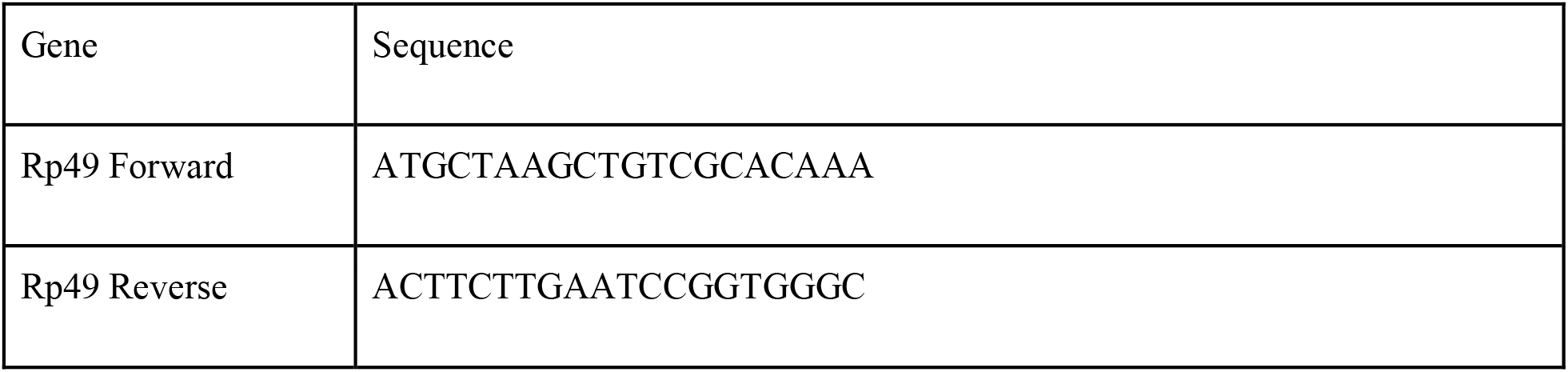
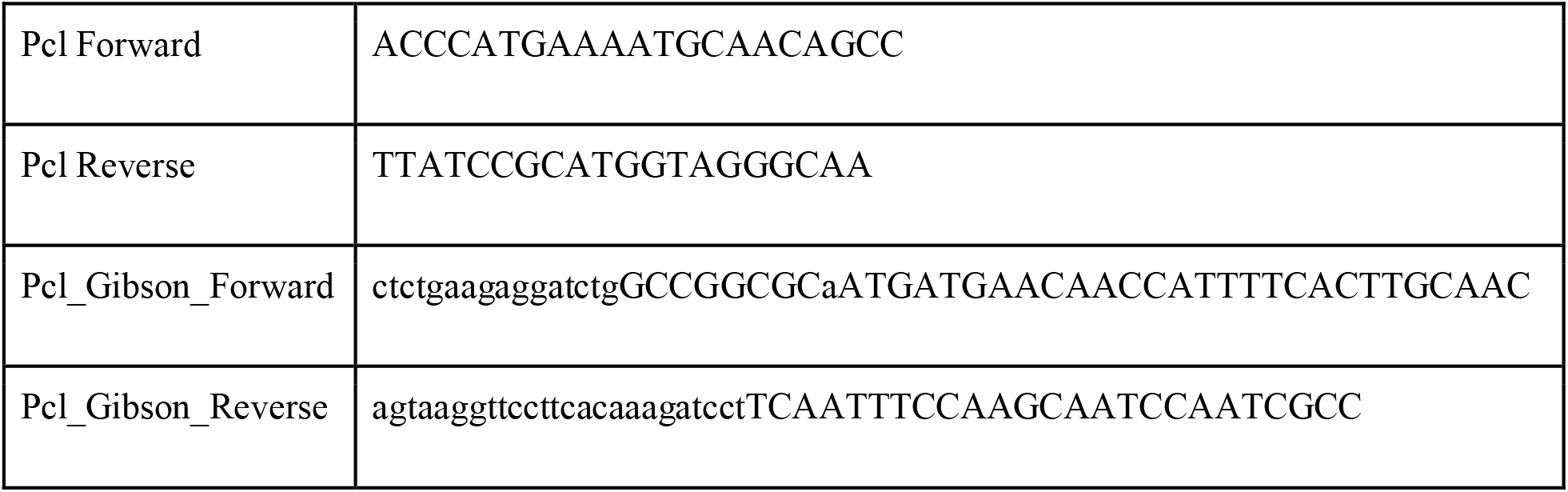

### Affinity purification of ribosome associated mRNA (TRAP)

300 heads (10,000 *Gr5a+* cells) per biological replicate were collected using pre-chilled sieves in liquid nitrogen on dry ice. Frozen heads were then lysed as previously described (*35*). From the lysate total RNA was extracted by TRIzol LS Reagent (ThermoFisher scientific, 10296010) as input. The remainder of the lysate was incubated with Dynabeads protein G (ThermoFisher scientific, 10004D) to preclear samples for 2 hours and subsequently incubated with Dynabeads protein G coated with an anti-Flag antibody (Sigma, F1804). The lysate-beads mixture was incubated at 4°C with rotation for 2 hours, then. RNA was extracted from ribosomes bound to the beads by TRIzol Reagent (*35*).

### Targeted DNA Adenine Methyltransferase Identification (TaDa)

For the *UAS-LT3-Dam::Pcl* construct, the coding region of the *Pcl* gene was amplified from the *pCRE-NDAM-Myc-DO69-Pcl* (gift from Bas Van Steensel, The Netherlands Cancer Institute) with primers listed above and assembled into the *UAS-LT3-DAM* plasmid (gift from Andrea Brand, University of Cambridge) using the NEBuilder® HiFi DNA Assembly kit based on manufacturer’s instructions (NEB). Transgenic animals were validated by RT-PCR for correct insert. These lines were crossed to *Gr5a-GAL4;tubulin-GAL80_ts_*. All animals were raised and maintained at 20 °C. Expression of *Dam::Pcl* and *Dam* were induced at 28 °C for 18-20 hours. For all experiments 300 heads of males and female flies were collected per replicate on dry ice by sieving. DNA was extracted from frozen heads following kit instructions (Invitrogen). For identification of methylated regions purified DNA was digested by DpnI followed by PCR purification of digested sequences. TaDa adaptors were ligated by T4 DNA ligase (NEB). Adapter ligated DNA was PCR amplified according to protocol (*28*), and subsequently purified. Purified DNA was digested with DpnII followed by sonication to yield fragments averaging 200-300bp. TaDa adaptors were removed from sonicated DNA by digestion. Sonicated DNA is used for library preparation (*28*).

### Library Preparation for TRAP and TaDa

RNA sequencing libraries were generated using the Ovation SoLo RNA-Seq System for *Drosophila* (Nugen, 0502-96). All reactions included integrated HL-dsDNase treatment (ArcticZymes, Cat. #70800-201). DNA sequencing libraries were generated using the Takara ThruPlex kit (cat #022818) using 3ng input and 10 cycles of PCR. All libraries were sequenced on the Illumina NextSeq platform (High-output kit v2 75 cycles) at the University of Michigan core facility.

### High Throughput RNA-seq Analysis

Fastq files were assessed for quality using FastQC (*74*). Reads with a quality score below 30 were discarded. Sequencing reads were aligned by STAR (*75*) to dmel-all-chromosomes of the dm6 genome downloaded from Ensemble, and gene counts were obtained by HTseq (*76*). Count files were used as input to call differential RNA abundance by DESeq2 (*77*). All pairwise comparisons were made to the control diet of the corresponding genotypes, such that sugar diet three days and sugar diet seven days were compared to the age matched control diet group. In *Pcl* mutant experiments, the pairwise comparison was made between sugar diet and control diet within the age-matched *Pcl_c429_* genotype group. A cutoff of *q*val<0.1 was used to call differentially expressed genes. Skew in log_2_fold changes was measured using the R package Skewness (e1071). RNAseq data visualization was carried out in R studio using ggplot2 and the following packages, pheatmap (*78*), Venneuler (*79*), and EnhancedVolcano (*80*). To cluster columns and rows in pheatmap “Ward.D ‘’ clustering was used.

### High Throughput TaDa and CATaDa Analysis

Fastq files were assessed for quality using FastQC (*74*). Reads with a quality score below 30 were discarded. The damidseq_pipeline was used to align, extend, and generate log2 ratio files (*Dam::Pcl/Dam*) in GATC resolution as described previously (*81*). In short, the pipeline uses Bowtie2 (*82*) to align reads to dmel-all-chromosomes of the dm6 genome downloaded from Ensemble, followed by read extension to 300 bp (or to the closest GATC, whichever is first). Bam output is used to generate the ratio file in bedgraph format. Bedgraph files were converted to bigwig and visualized in the UCSD genome browser. Correlation coefficients and PCA plot between biological replicates were computed by multibigwigSummary and plotCorrelation in deepTools (*83*). Fold Change traces for SD/CD of log_2_(*Dam::Pcl/Dam*) were generated by calculating the mean coverage profile of all replicates for each condition and subsequently calculating fold change between the sugar diet and control diet condition with deepTools bigwigCompare (*83*). Peaks were identified from log_2_(*Dam::Pcl/Dam*) ratio files using find_peaks (FDR<0.01) (*81*). To do this, the binding intensity thresholds are identified from the dataset, the dataset is then shuffled randomly, and the frequency of consecutive regions (i.e. GATC fragments or bins) with a score greater than the threshold is calculated. The FDR is the observed / expected for a number of consecutive fragments above a given threshold. Association of genes to peaks was made using the peaks2genes script (*81*) and dm6 genome annotations. Overlapping intervals or nearby intervals were merged into a single interval using MergeBED in Bedtools (*84*). Intervals common in all replicate peak files were identified by Multiple Intersect in Bedtools (*84*). DiffBind was used to determine differentially bound sites on peak files based on differences in read intensities (*85*).

For CATaDa experiments, all analyses were performed similar to those of TaDa with the exceptions that 1) *Dam* only profiles were not normalized as ratios but shown as non-normalized binding profiles, 2) *Dam* only coverage plots were generated by converting bam files to bigwig files normalized to 1x dm6 genome, and 3) peaks were called using MACS2 call peaks on alignment files without building the shifting model with an of FDR<0.05 (*86*).

To determine the proportion of genes that fit within the various chromatin domain subtypes, we first matched *Dam::Pcl/Dam* targets to coordinates identified by Filion et al and then determined their gene count in each chromatin subtype (observed) compared to the whole genome (expected).

### iPAGE Analysis

All Gene Ontology (GO) term enrichment analysis was performed using the iPAGE package (*34*), using gene-GO term associations extracted from the Flybase dmel 6.08 2015_05 release (*87*). For analysis of TRAP data, iPAGE was run in continuous mode, with log_2_fold changes divided into seven equally populated bins. For analysis of TaDa and TF regulatory targets, iPAGE was run in discrete mode, using the groups specified for each calculation. For all discrete calculations, independence filtering was deactivated due to the less informative available signal. All other iPAGE settings were left at their default values. All shown GO terms pass the significance tests for overall information described in (*34*); in addition, for each term, individual bins showing especially strong contributions (*p*-value such that a Benjamini-Hochberg FDR (*88*) calculated across that row, yields *q*<0.05) are highlighted with a strong black box.

### PREdictor

Identification of predicted PRE sites was performed exactly as described in (*33*), using the dm6 genome. As suggested in (*33*), we use a threshold confidence score of 0.8 to identify the PREs used in the present analysis. A complete list of predicted PREs, with accompanying confidence scores, is shown in Supplementary File 1. Enrichments of overlap between different PRE classes and *Pcl* occupancy locations were calculated by comparing the observed overlap frequency with the overlaps for 1,000 random shuffling of the binding/differential binding peak locations (calculated using bedtools 2.17.0 (*84*)).

### Calculation of regulatory targets of transcription factors

To identify likely targets of each transcription factor of interest, we drew upon the transcription factor binding site calculations described in (*33*), in which the motif of each transcription factor was scanned along every base pair of the *D. melanogaster* dm6 genome using FIMO (*89*), and the base pair-wise binding results converted to robust z-scores. For each TF, we then considered its regulon to consist of all genes with at least one binding site with z-score above a TF-specific threshold within 2 kb upstream of the beginning of the gene. We identified TF-specific thresholds by manual inspection of a plot of the average expression changes between conditions vs. threshold, aiming to identify a point of maximum information content relative to noise (see Supplemental Figure 9 for the plots used to identify TF-specific z-score thresholds). Once the set of targets (regulon) for each factor was identified, we tested for significant enrichment or depletion of overlaps between the regulons using Fisher’s exact test, reporting Benjamini-Hochberg false discovery rates (FDRs) (*88*). All calculated odds ratios were positive, indicating enriched overlaps between the regulons.

### STRING network analysis

To develop a functional network between the candidate neural targets of cad, Ptx1, nub, GATAe and Scro, we uploaded genes categorized into neural/signaling GO terms based on DAVID functional annotations (*90, 91*) to the STRING database search (*37*). Genes were clustered by their reported protein-protein interactions and corresponding confidence scores (*37*) and plotted in Cytoscape (v3.7.1) (*92*). In this network edges do not represent direct protein-protein interaction but rather represent a functional interaction. For network see Supplementary File 6.

### Data Analysis and Statistics

Statistical tests, sample size, and *p* or *q* values are listed in each figure legend. Data were evaluated for normality and appropriate statistical tests applied if data were not normally distributed, all the tests, biological samples, and the *p* and *q* values are listed in the figure legends and specific analysis under each methods session. Because the inferential value of a failure to reject the null hypothesis in frequentist statistical approaches is limited, for all RNA-seq expression datasets, we coupled our standard differential expression with a test for whether each gene could be flagged as ‘significantly not different’. Defining a region of practical equivalence (ROPE) as a change of no more than 1.5-fold in either direction, we tested the null hypothesis of at least a 1.5-fold change for each gene, using the gene-wise estimates of the standard error in log_2_fold change (reported by Deseq2) and the assumption that the actual log_2_fold changes are normally distributed. Rejection of the null hypothesis in this test is taken as positive evidence that the gene’s expression is not changed substantially between the conditions of interest. Python code for the practical equivalence test can be found on Github as calc_sig_unchanged.py. All data in the figures are shown as Mean ± SEM, **** *p* < 0.0001, *** *p* < 0.001, ** *p* < 0.01, \**p*< 0.05 unless otherwise indicated.

## Acknowledgements

We thank Sundeep Kalantry, Josie Clowney, and Giacomo Cavalli for helpful comments and discussions. We also thank the University of Indiana at Bloomington, the VDRC, the FLYORF stock collections, and all the investigators who kindly shared fly lines with us. We also thank Daniel Wilinski for help with data analysis, Dion Dickman and Xun Chen for helpful comments on riboTRAP, and Margarita Brovkina (E.J. Clowney lab) for helpful discussions on TaDa analysis. Matt Burkhard assisted with analysis of expression changes for different Gene Ontology terms. Julia Kuhn designed some of the graphics for the manuscript. This work was funded by NIH R00 DK-97141 and NIH 1DP2DK-113750, the Klingenstein-Simons Fellowship in the Neurosciences, and the Rita Allen Foundation (to M.D.) and NIH R35 GM-128637 (to P.L.F).

## Author contributions

A.V performed all experiments and analyzed RNAseq, TaDa, and CATaDa datasets, with the exception of *in vivo* calcium imaging. B.T.G helped with PER and triglyceride measurements. C.E.M performed *in vivo* calcium imaging. M.K and P.L.F developed the PREdictor, calculated regulatory targets of transcription factors, ran and analyzed Gene Ontology term enrichment analysis, provided statistical consultation, and tested for practical significance. M.D oversought the project and secured funding. A.V and M.D designed the experiments, wrote the manuscript, and prepared the figures with input from all authors.

## Data availability

All high throughput sequencing data files can be found on Gene Expression Omnibus GSE146245.

## Supplementary Figures

**Fig. S1.**
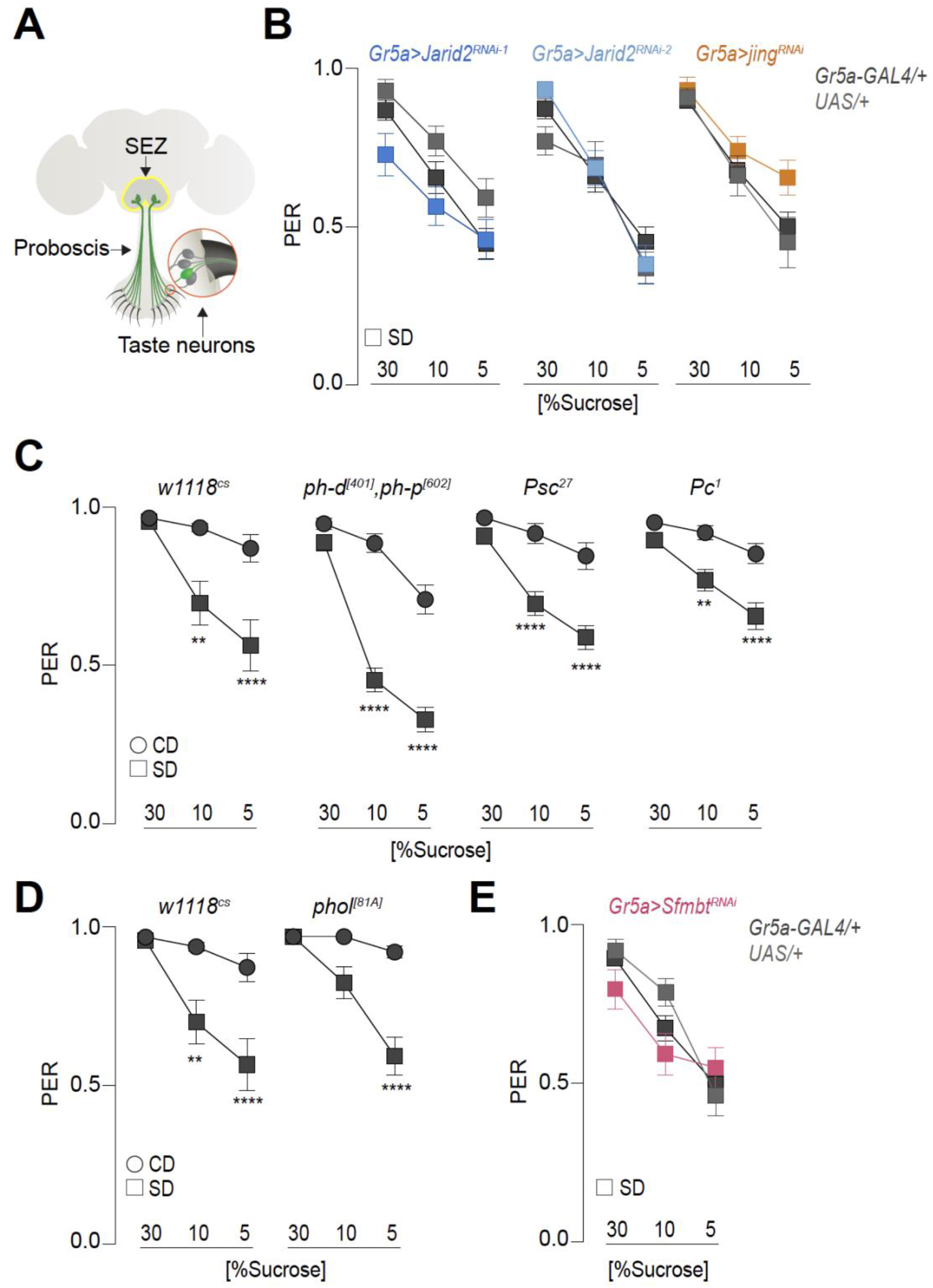
The PRC1 and PhoRC complex are not required for sugar diet mediated sweet taste defects. **(A)** The fly proboscis contains the sweet sensory neurons soma (green) with dendrites extending into the hair sensilla and their axons projecting onto the subesophageal zone (SEZ, delimited in yellow) of the fly brain. **(B-E)** Taste responses (y axis) to stimulation of the labellum with 30%, 10% and 5% sucrose (x axis) of age-matched males of **(B)** *Gr5a>Jarid2_RNAi-1_* (dark blue), *Gr5a>Jarid_RNAi-2_* (light blue), and *Gr5a>jing_RNAi-1_* (orange) and parental transgenic control (gray, crossed to *w1118_cs_*) flies on a sugar diet for 7 days. n = 25-34, Kruskal-Wallis Dunn’s multiple comparisons, comparisons to transgenic controls. **(C)** *w1118_cs_*, *ph-d_[401]/+_ph-p_[602]/+_*, *Psc_27/+_*,and *Pc_1/+_* flies on a control (circle) or sugar (square) diet for 7 days. n = 30-42, Kruskal-Wallis Dunn’s multiple comparisons, comparisons to control diet within each genotype group. **(D)** *w1118_cs_*, *phol_[81A]/+_* flies on a control (circle) or sugar (square) diet for 7 days. n = 32-34, Kruskal-Wallis Dunn’s multiple comparisons, comparisons to control diet within each genotype group. **(E)** *Gr5a>Sfmbt_RNAi_* (dark pink) and parental transgenic control (gray, crossed to *w1118_cs_*) flies on a sugar diet for 7 days. n = 26-46, Kruskal-Wallis Dunn’s multiple comparisons, comparisons to transgenic controls. All data shown as mean ± SEM, ∗∗∗∗*p* < 0.0001, ∗∗∗*p* < 0.001, ∗∗*p* < 0.01, and ∗*p* < 0.05 for all panels unless indicated.

**Fig. S2.**
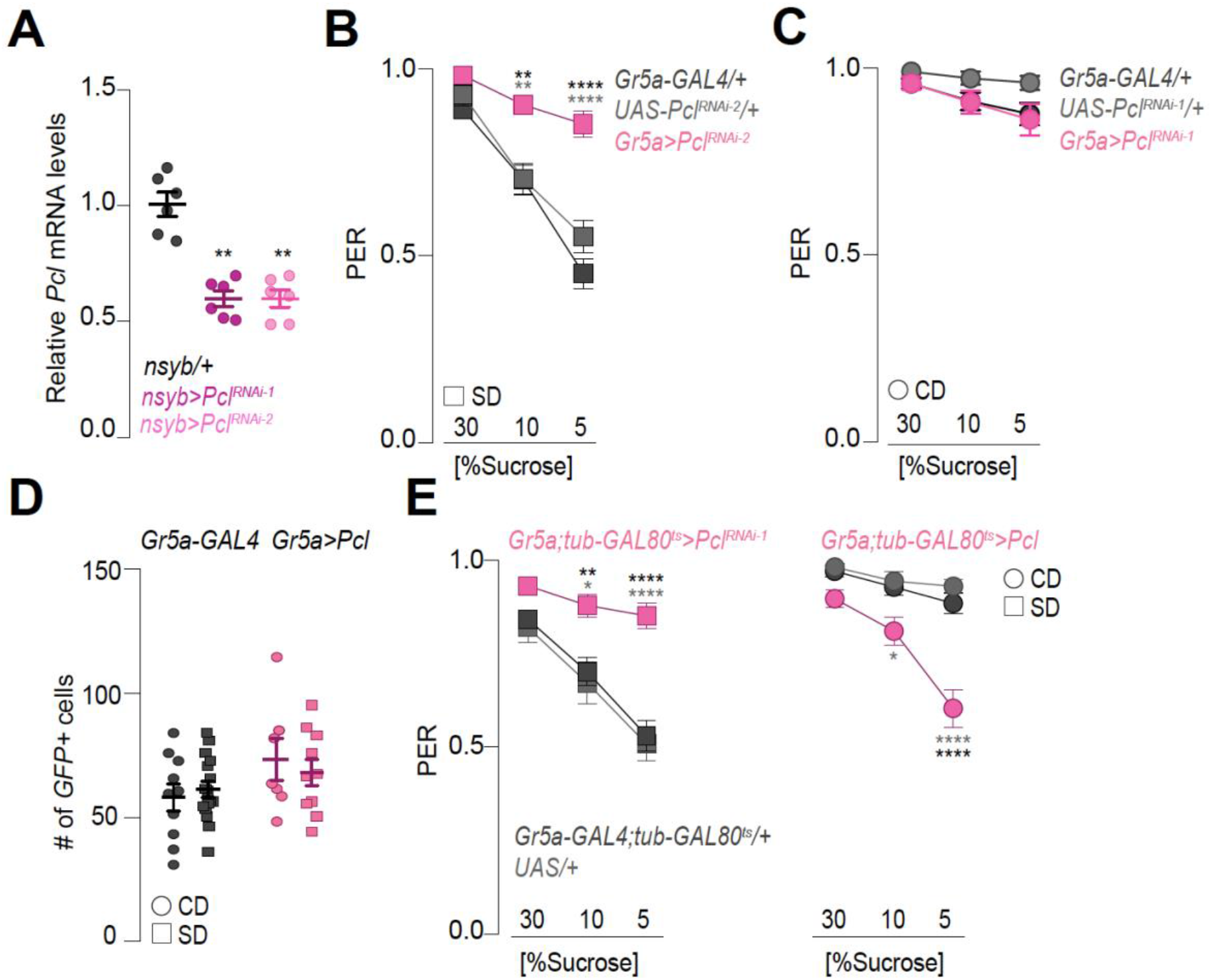
The effects of *Pcl* on sweet taste are independent of changes in cell number or development. **(A)** Fold change of *Pcl* mRNA levels in the heads of flies with and without pan-neuronal (*nsyb-GAL4*) *Pcl* knockdown with two independent RNAi lines (dark gray, plum, and pink respectively) measured by qPCR and normalized to the gene *Rp49*. n=6, Kruskal-Wallis Dunn’s multiple comparisons, compared to GAL4 transgenic control (crossed to *w1118_cs_*). **(B-C)** Taste responses (y axis) to stimulation of the labellum with 30%, 10% and 5% sucrose (x axis) of age-matched males of **(B)** *Gr5a>Pcl_RNAi-2_* and parental transgenic control (gray, crossed to *w1118_cs_*) flies on a sugar diet for 7 days. n = 42-51, Kruskal-Wallis Dunn’s multiple comparisons, comparisons to transgenic controls. **(C)** *Gr5a>Pcl_RNAi-1_* and parental transgenic control (gray, crossed to *w1118_cs_*) flies on a control diet for 7 days. n =30-35, Kruskal-Wallis Dunn’s multiple comparisons, comparisons to transgenic controls. **(D)** Quantification of the number of sweet taste GFP-labeled cells in the labella of *Gr64f;CD8-GFP* flies crossed to *w1118_cs_* (as control, gray) or *Gr64f;CD8-GFP>Pcl (pink)* on a control (circle) or sugar (square) diet for 7 days. n=5-16 probosces, no significance, Kruskal-Wallis Dunn’s multiple comparisons, comparison to control diet of each genotype. **(E)** Taste responses (y axis) to stimulation of the labellum with 30%, 10% and 5% sucrose (x axis) of age-matched males of *Gr5a;tubulin-GAL80_ts_>Pcl_RNAi-1_*, *Gr5a;tubulin-GAL80_ts_>Pcl,* and parental transgenic control (gray, crossed to *w1118_cs_*) flies on a control (circle, right) or sugar (square, left) diet for 7 days. n = 34-49, Kruskal-Wallis Dunn’s multiple comparisons, comparisons to transgenic control. All data shown as mean ± SEM, ∗∗∗∗*p* < 0.0001, ∗∗∗*p* < 0.001, ∗∗*p* < 0.01, and ∗*p* < 0.05 for all panels unless indicated.

**Fig. S3.**
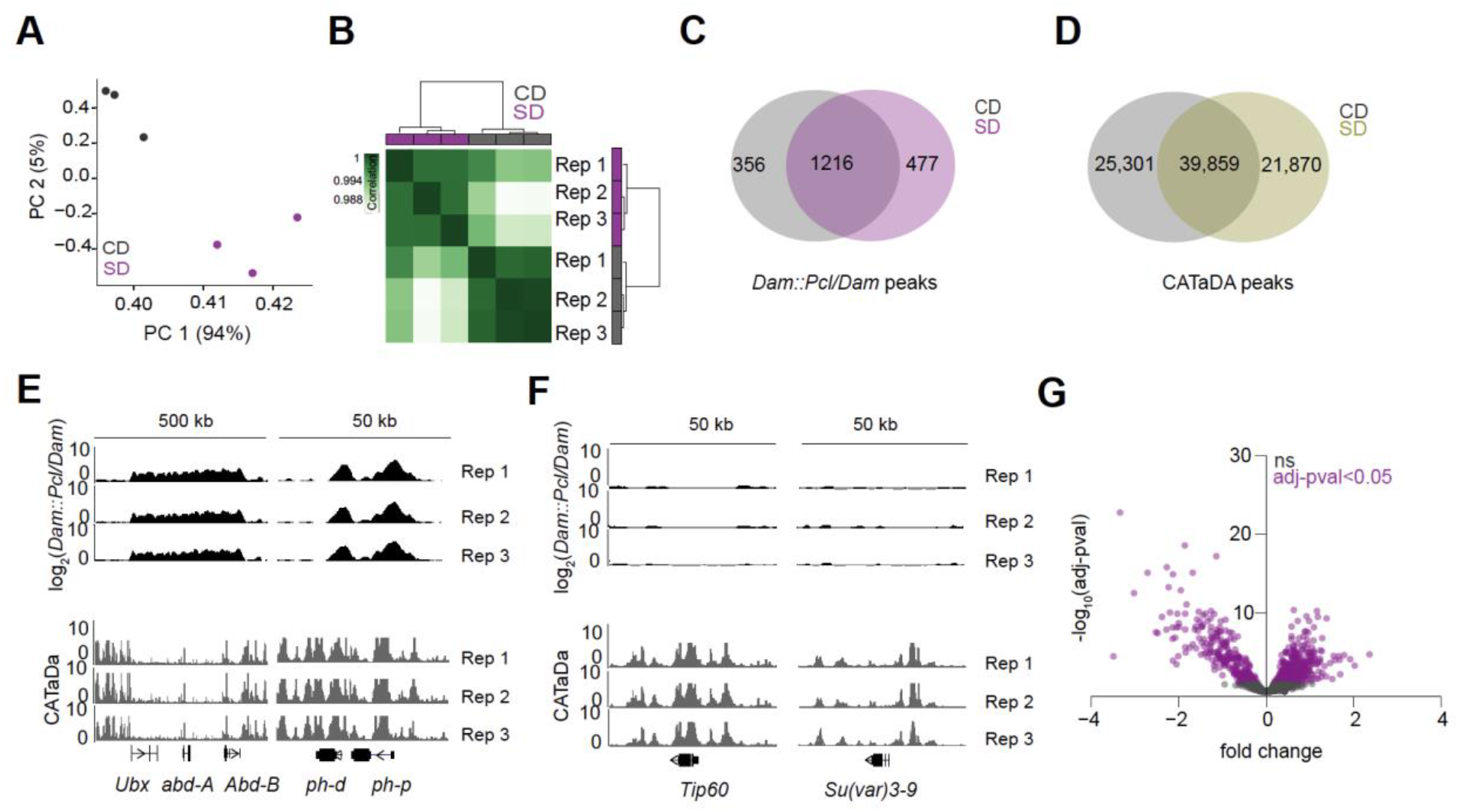
Chromatin occupancy and accessibility analysis of *Pcl* in the *Gr5a+* neurons. **(A)** Principal component analysis of normalized log_2_(*Dam::Pcl/Dam*) flies on a control (CD, gray) or sugar (SD, purple) diet. **(B)** The three biological replicates of log_2_(*Dam::Pcl/Dam*) on a control (CD, gray) or sugar (SD, purple) diet are shown, with high Pearson correlation coefficients within each dietary condition, and low correlation coefficients between dietary conditions. Scale shows degree of correlation. **(C-D)** Left panel: overlap of log_2_(*Dam::Pcl/Dam)* chromatin binding peaks between a control (gray) and sugar (purple) diet (find_peaks, *q*<0.01). Right panel: overlap of CATaDa chromatin peaks between a control (gray) and sugar (yellow) diet (MACS2, *q*<0.05). **(E-F)** Log_2_*(Dam::Pcl/Dam*) chromatin binding coverage at *Ultrabithorax* (*Ubx*), *abdominal A* (*abd-A*), *Abdominal B* (*Abd-B*), *polyhomeotic distal* (*ph-d*) and *polyhomeotic proximal* (*ph-p*) (Top panel, E), and *Tat interactive protein 60* (*Tip60*) and *Suppressor of variegation 3-9* (*Su(var)3-9*) (Top panel, F). Bottom panel: CATaDa chromatin coverage (chromatin accessibility) of the three biological replicates at the loci mentioned above. **(G)** Volcano plot showing the differential chromatin binding of *Dam::Pcl* on a sugar diet compared to a control diet, (Wald test, *q*<0.05, purple), non-significant peaks are shown in black (ns).

**Fig. S4.**
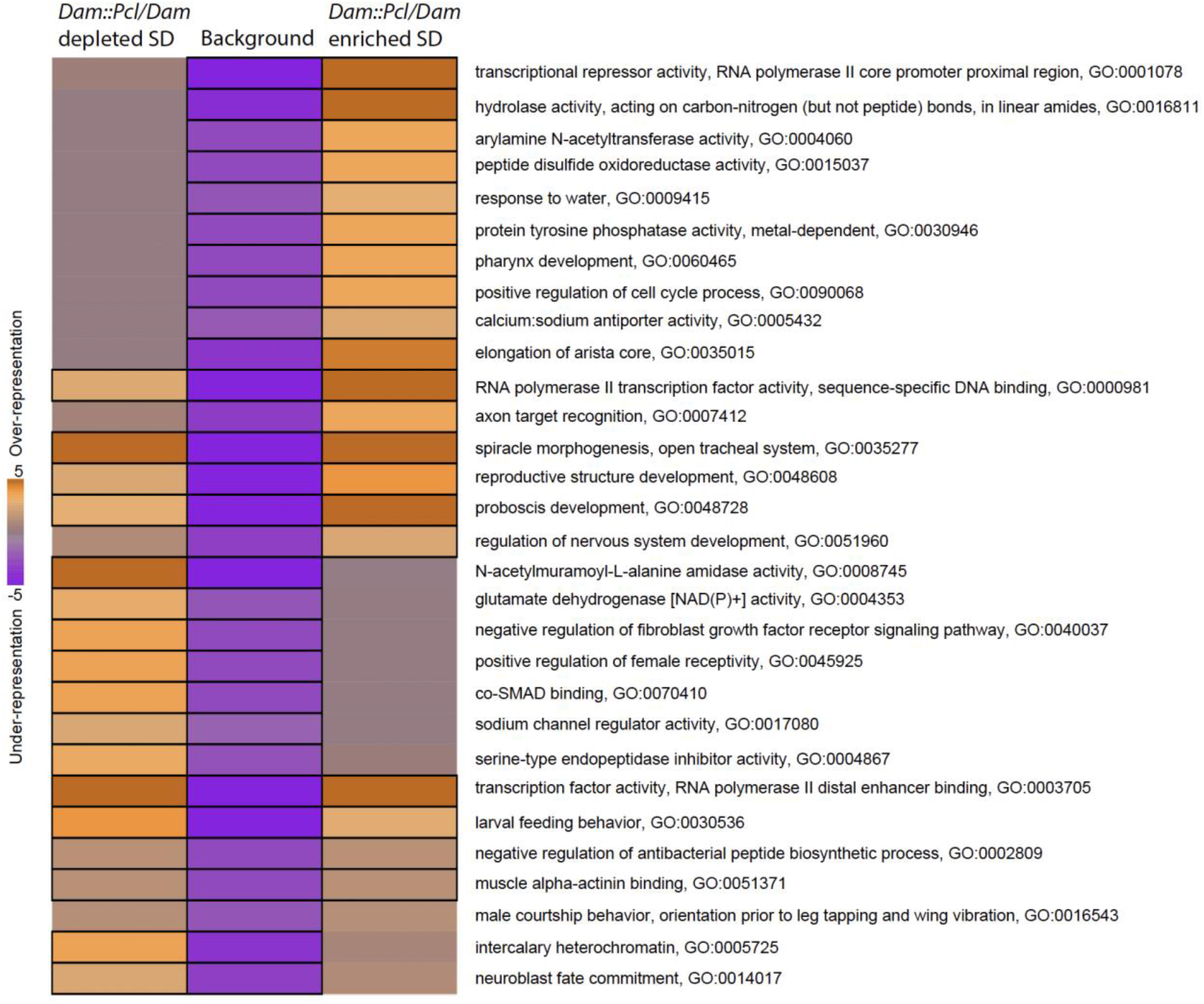
Pathway enrichment analysis of *Pcl* chromatin targets in the *Gr5a+* neurons. iPAGE identification of pathways depleted (left) or enriched (right) compared to background gene list (middle) from the differentially bound chromatin peaks by *Dam::Pcl* flies on a sugar diet compared to a control diet. Scale represents over-representation (orange) or under-representation (purple) of genes within a specific bin for the corresponding GO term. Black outlined boxes represent *q*<0.05.

**Fig. S5.**
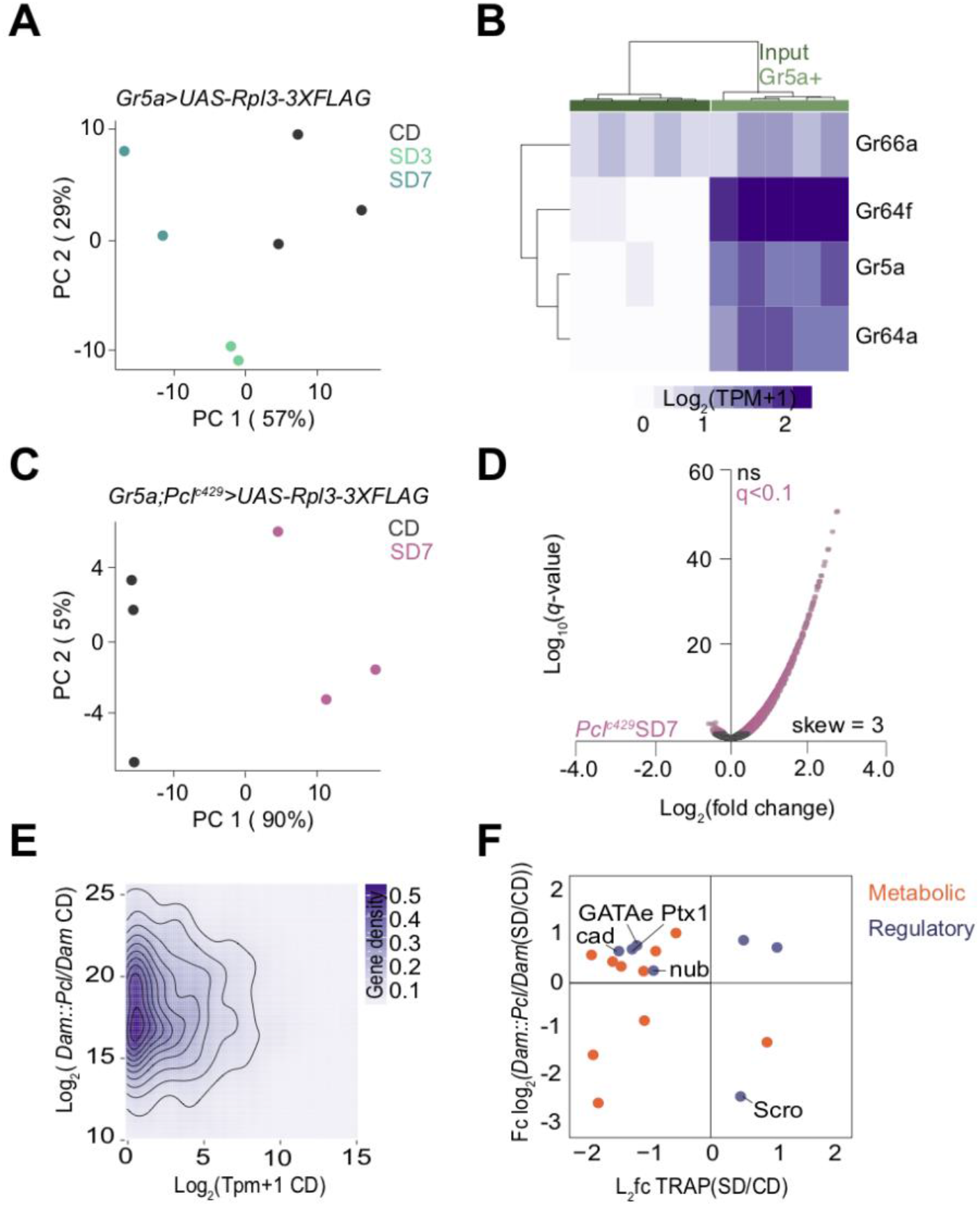
PRC2.1 mediates the transcriptional responses of the *Gr5a+* neurons to diet. **(A)** Principal component analysis of age matched male *Gr5a>UAS-Rpl3-3XFLAG* flies on a control diet (CD, gray), sugar diet for three days (SD3, mint), and sugar diet for seven days (SD7, teal). **(B)** Normalized read counts for *Gustatory receptor Gr66a* (bitter), *Gustatory receptor Gr64f*, *Gustatory receptor Gr5a*, and *Gustatory receptor Gr64a* (all sweet) from the input fraction (dark green box) and *Gr5a+* fraction (light green box) specific conditions. Both samples also include the dietary conditions (not specified in figure), hues of purple from light to dark represent low to high expression. **(C)** Principal component analysis of age matched male *Gr5a;Pcl_c429_>UAS-Rpl3-3XFLAG* flies on a control diet (CD, gray) and sugar diet for seven days (SD7, pink). **(D)** Volcano plot with changes in gene expression in the *Gr5a+* neurons of age matched male *Gr5a;Pcl_c429_>UAS-Rpl3-3XFLAG* flies on a SD for 7 days (*Pcl_c429_*SD7, pink) compared to the control diet condition in the same genotype (Wald test, *q*<0.1), n=3 replicates per condition (∼10,000 *Gr5a+* cells per replicate). **(E)** Normalized reads (Tpm+1) from TRAP for genes (x-axis) bound by *Pcl* and normalized to log_2_(*Dam::Pcl/Dam*) reads at these genes (y-axis) on a control diet. Scale represents gene density (light to dark purple). **(F)** Fold change of *Dam::Pcl* chromatin binding for genes differentially bound on sugar diet compared to control diet (y-axis) (*q*<0.05) and their respective differentially expressed log_2_ fold changes on a sugar diet compared to control diet (x-axis) (*q*<0.1). Genes are shown as circles and colored based on GO term category, metabolism (orange) and regulatory (lavender).

**Fig. S6.**
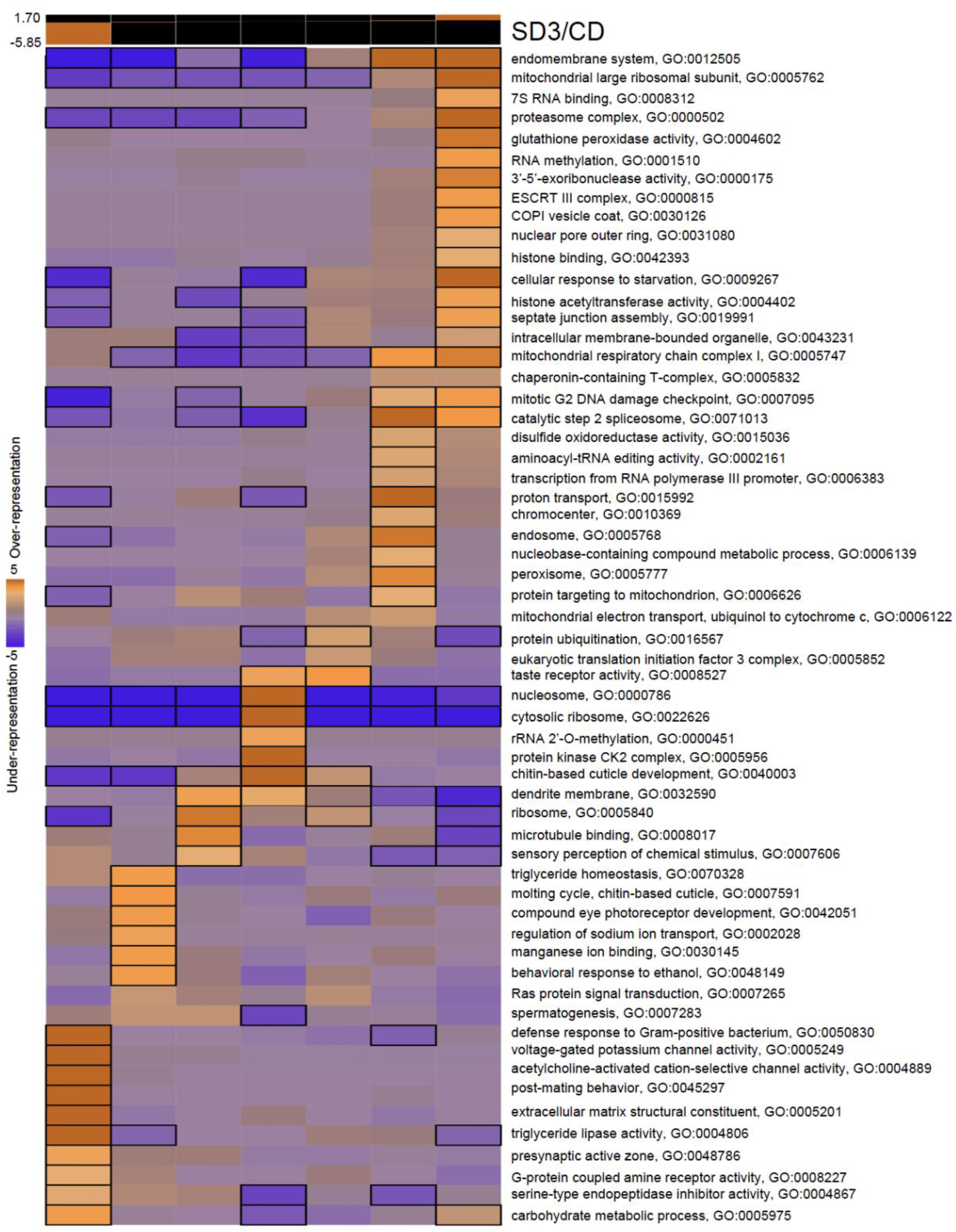
Pathway enrichment analysis of the *Gr5a+* neurons in flies fed a sugar diet for 3 days. iPAGE identification of pathways enriched in age-matched male *Gr5a>UAS-Rpl3-3XFLAG* flies on a sugar diet for three days compared to a control diet. Bins (top) show the range of log_2_ fold changes for genes within their corresponding GO terms. Scale represents over-representation (orange) or under-representation (purple) of genes within a specific bin for the corresponding GO term. Black outlined boxes represent *q*<0.05.

**Fig. S7.**
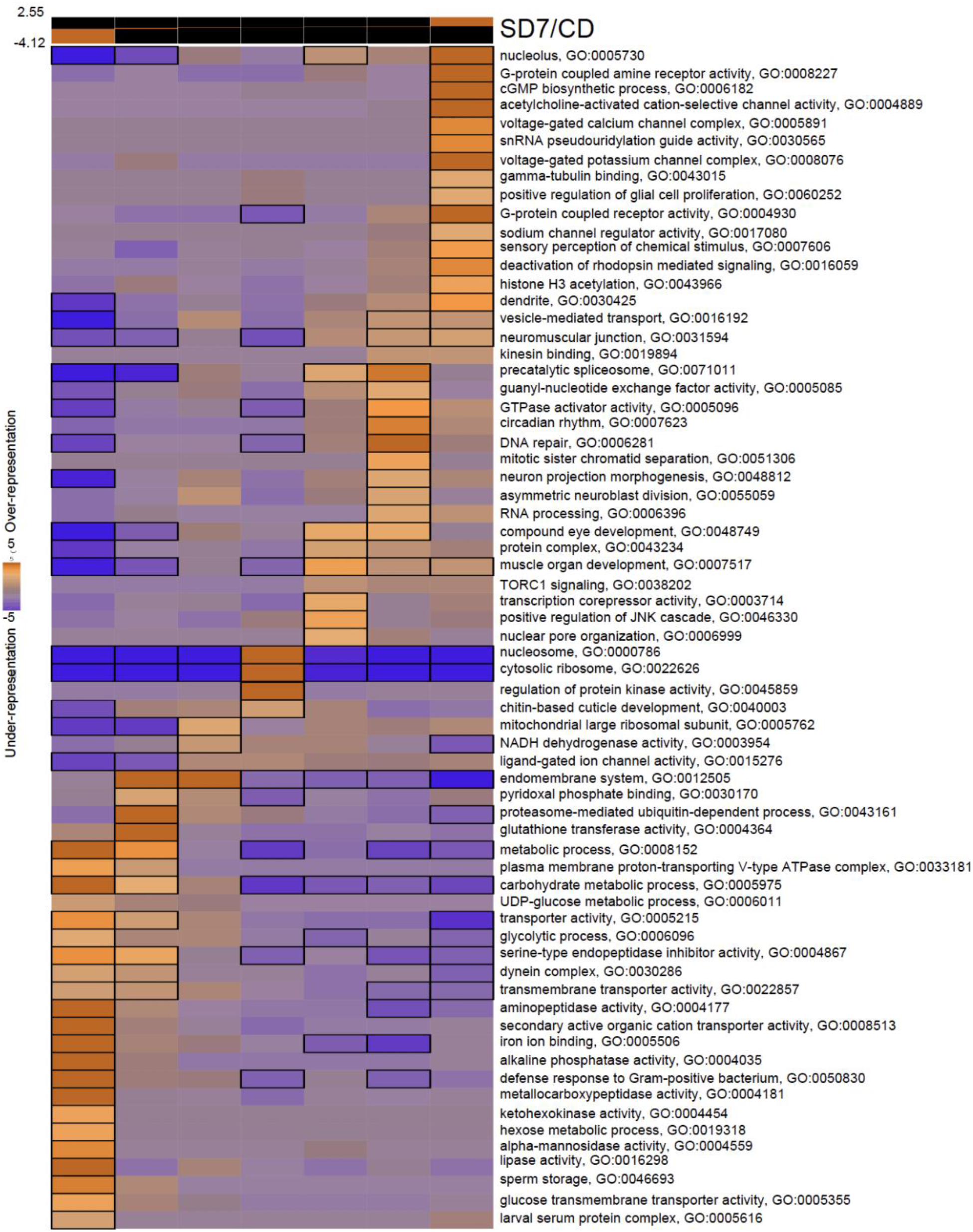
Pathway enrichment analysis of the *Gr5a+* neurons in flies fed a high sugar diet for 7 days. iPAGE identification of pathways enriched in age-matched male *Gr5a>UAS-Rpl3-3XFLAG* flies on a sugar diet for seven days compared to a control diet. Bins (top) show the range of log_2_ fold changes for genes within their corresponding GO terms. Scale represents over-representation (orange) or under-representation (purple) of genes within a specific bin for the corresponding GO term. Black outlined boxes represent *q*<0.05.

**Fig. S8.**
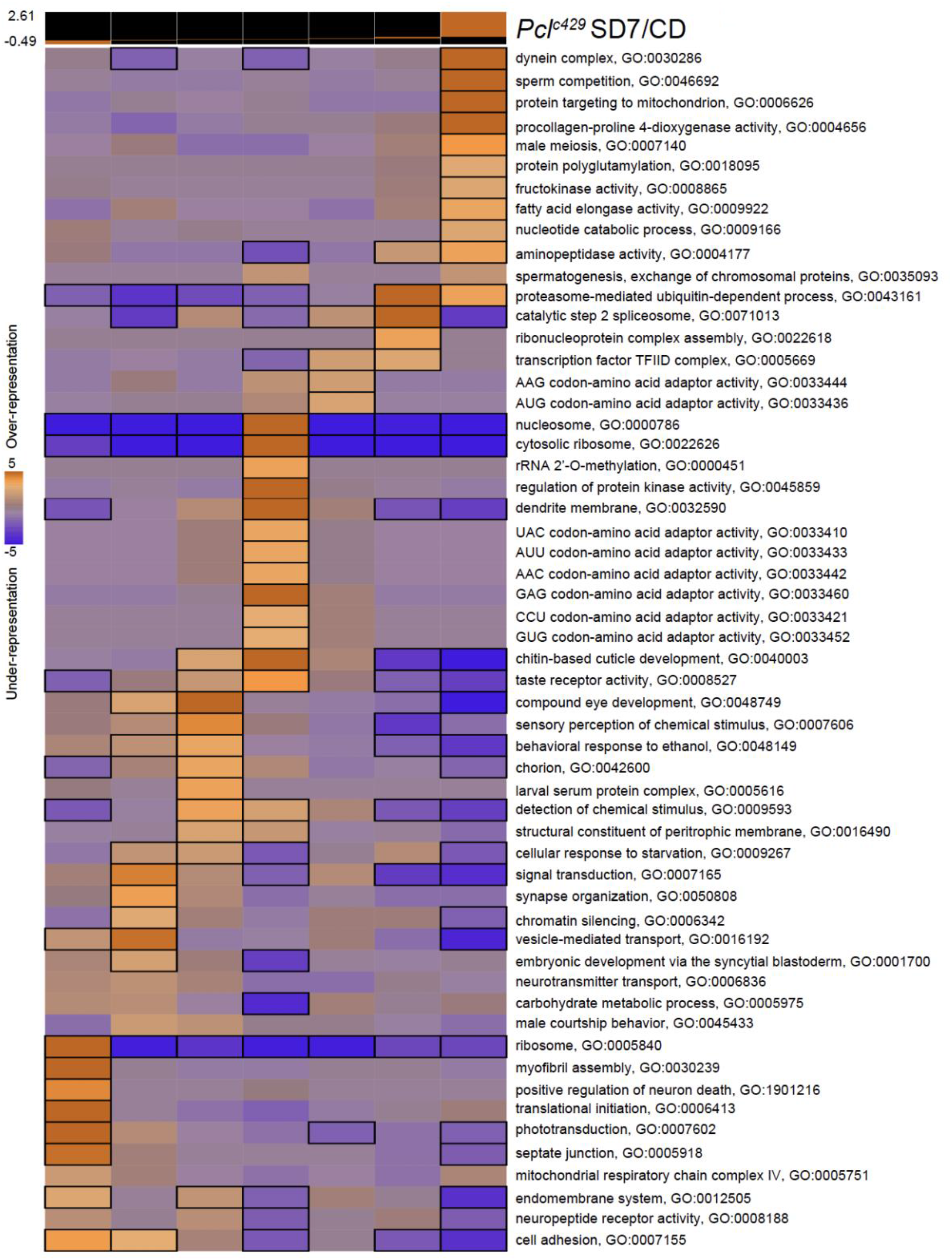
Pathway enrichment analysis of the *Gr5a+* neurons in *Pcl_c429_* mutant flies fed a high sugar diet for 7 days. iPAGE identification of pathways enriched in age-matched male *Gr5a;Pcl_c429_>UAS-Rpl3-3XFLAG* flies on a sugar diet for seven days compared to a control diet. Bins (top) show the range of log_2_ fold changes for genes within their corresponding GO terms. Scale represents over-representation (orange) or under-representation (purple) of genes within a specific bin for the corresponding GO term. Black outlined boxes represent *q*<0.05.

**Fig. S9.**
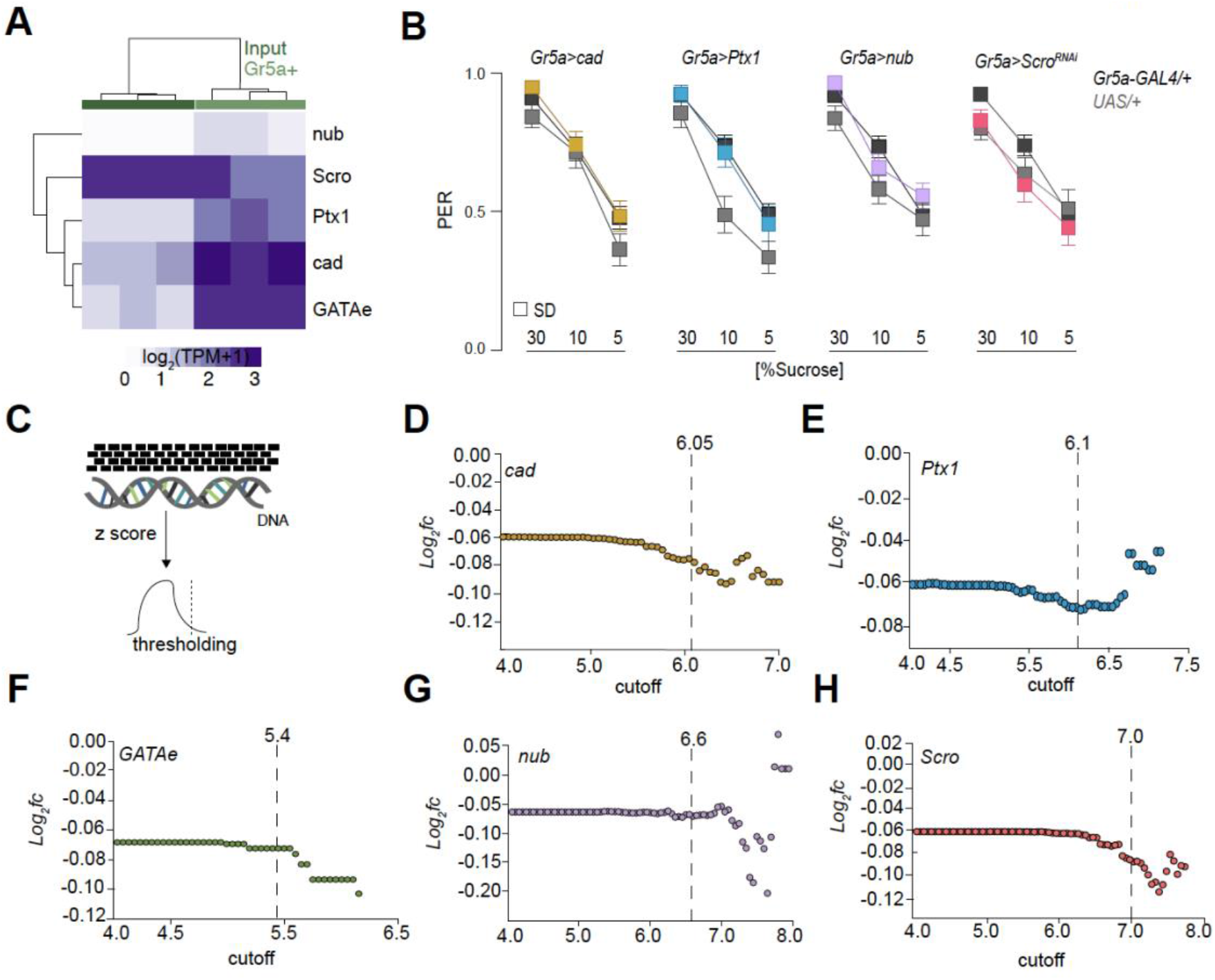
Identification of the transcriptional program mediated by PRC2.1 on a sugar diet. **(A)** Normalized read counts for *Scro*, *cad*, *nub*, *GATAe*, and *Ptx1* in the input fraction (dark green box) and the *Gr5a+* fraction (light green box) from *Gr5a>UAS-Rpl3-3XFLAG* age-matched male flies on a control diet; hues of purple from light to dark show low to high expression. **(B)** Taste responses (y axis) to stimulation of the labellum with 30%, 10% and 5% sucrose (x axis) of age-matched males of *Gr5a>cad (gold), Gr5a>Ptx1 (blue), Gr5a>nub (lavender), Gr5a>Scro_RNAi_* (coral), and parental transgenic control (gray, crossed to *w1118_cs_*) flies on a sugar diet for 7 days. n = 31-51, Kruskal-Wallis Dunn’s multiple comparisons, comparisons to transgenic controls. **(C)** Schematic of how the targets of the 5 transcription factors in Figure 5 were identified: the DNA binding motifs for each transcription factor were scanned along every base pair of the *D. melanogaster* genome using FIMO, and the basepair-wise binding results converted to robust z-scores. **(D-H)** Threshold scores used to identify the regulons of each transcription factor. Y-axis represents the averaged log_2_ fold change of the predicted targets based on the cutoff score, respectively.

**Fig. S10.**
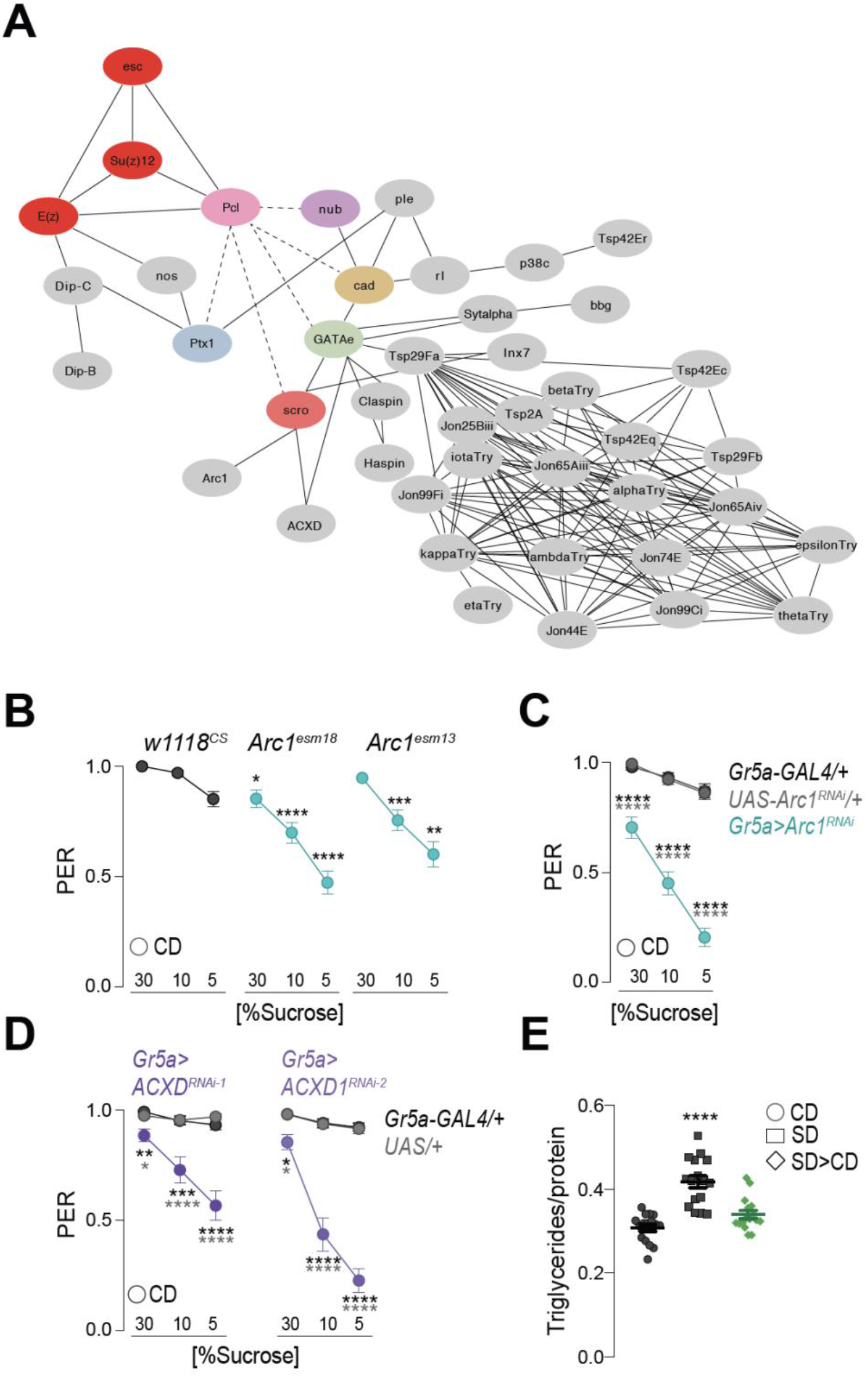
PRC2.1 modulates the synaptic and neural properties of the *Gr5a+* neurons on a sugar diet. **(A)** Functional network created with STRING featuring PRC2.1, *cad*, *Ptx1*, *GATAe*, *nub*, *Scro,* and their predicted targets (gray) from Figure 5C that can be classified into the GO term category neural/signaling. Each node represents a gene. Edges indicate functional protein-protein interactions, not direct interactions. Edges represented with dashed lines are interactions identified in this study. **(B-D)** Taste responses (y axis) to stimulation of the labellum with 30%, 10% and 5% sucrose (x axis) of age-matched males of **(B)** *w1118_cs_*, *Arc1_esm18_*, and *Arc1_esm13_* flies on a control diet for 7 days. n = 38-45, Kruskal-Wallis Dunn’s multiple comparisons, comparisons to transgenic controls. **(C)** *Gr5a>Arc1_RNAi_ (turquoise),* and parental transgenic control (gray, crossed to *w1118_cs_*) flies on a control diet for 7 days. n = 28-47, Kruskal-Wallis Dunn’s multiple comparisons, comparison to control diet. **(D)** *Gr5a>ACXD_RNAi-1_* (dark purple) *and Gr5a>ACXD_RNAi-2_* (light purple), and parental transgenic controls (gray, crossed to *w1118_cs_*) flies on a control diet for 7 days. n = 28-47, Kruskal-Wallis Dunn’s multiple comparisons, comparison to control diet. **(E)** Triglyceride levels normalized to protein in *w1118_CS_* flies fed a control (gray, circle) or sugar (gray, square) diet for 14 days, and flies fed a SD>CD reversal diet for a total of 14 days (7 days in each diet). n=16, two-way ANOVA with Sidak’s multiple comparisons test, comparisons to control diet of each genotype. All data are shown as mean ± SEM, ∗∗∗∗*p* < 0.0001, ∗∗∗*p* < 0.001, ∗∗*p* < 0.01, and ∗*p* < 0.05 for all panels unless indicated.

## Supplementary Files

**Supplementary file 1:**

PRE prediction calls for the dmel6.08 genome. Relating to Figure 3.

**Supplementary file 2:**

Differential binding analysis of *Dam::Pcl/Dam* on a sugar diet compared to control diet.

**Supplementary file 3:**

Differential expression analysis of TRAP experiments for sugar diet 3 (SD3), sugar diet 7 (SD7), *Pcl_c429_* SD7 (all comparisons to control diet) and CD>CD and SD>CD.

**Supplementary file 4:**

Transcription factor regulatory target calls relating to Figures 5.

**Supplementary file 5:**

iPAGE identification of pathways over-represented or under-represented in the predicted gene targets of *GATAe, cad, nub, Ptx1* and *Scro*.

**Supplementary file 6:**

STRING interaction network relating to Supplement Figure 14.

**Supplementary file 7:**

Gene table for heatmap relating to Figure 6D.

